# Single-cell, clonal and spatial atlases of cranial placodes illuminate their specification and evolution

**DOI:** 10.64898/2026.04.01.715621

**Authors:** Aliia Murtazina, Yuliia Fatieieva, Felix Waern, Helen R. Maunsell, Ankita Thawani, Bettina Semsch, Johan Boström, Caleb C. Reagor, Polina Kameneva, Karina Araslanova, Sergey Isaev, Franziska Schelb, Kaj Fried, Alek G. Erickson, Alexander Klimovich, Andrea Streit, Lena M. Kutscher, Iryna Kozmikova, Zbynek Kozmik, Emma R. Andersson, Gerhard Schlosser, Andrew K. Groves, Igor Adameyko

## Abstract

The vertebrate head is defined by complex sensory structures derived from cranial placodes. Placodes arise alongside the neural crest at the neural plate border, yet the mechanisms governing their identity, diversification, and evolutionary origins are unclear. We present an integrated single-cell, spatial, and clonal atlas of placode development to resolve the dynamics of their lineage segregation. Combining single-cell RNA-sequencing, spatial transcriptomics, and high-resolution clonal tracing, we show that placodal and neighboring progenitors form a continuous transcriptional landscape with gradual transitions between domains. Domain boundary cells co-express markers of adjacent territories, suggesting transient bipotent states. Consistent with this, clonal analysis reveals sharing of progenitors between neighboring placodes, supporting a model of competitive segregation. Comparisons with amphioxus suggests that vertebrate olfactory placodes emerged from an ancestral neuroectoderm that later partitioned into distinct neural and olfactory domains. Our findings provide a unified framework for understanding the developmental and evolutionary origins of vertebrate sensory organs.

## Introduction

The cranial placodes, together with the neural crest, represent defining innovations of the vertebrate lineage^1–3^. These ectodermal structures generate the peripheral neurons and sensory structures of the head, underpinning the elaborate cranial sensory apparatus that distinguishes vertebrates from their invertebrate relatives^1,4,5^. Each neurogenic placode contributes to specific sensory organs and ganglia. The olfactory placode forms the olfactory epithelium and olfactory receptor neurons, which project to the forebrain’s olfactory bulbs^6^. The lens placode invaginates to produce the lens of the eye, while the adjacent optic cup, derived from the neural tube, forms the retina - together establishing the vertebrate visual system^7^. The adenohypophyseal placode contributes to the anterior lobe of the pituitary gland (adenohypophysis), a key endocrine organ^8^. The trigeminal placode, located at an intermediate position along the anterior-posterior axis of the developing head, generates the somatosensory neurons of the trigeminal ganglion, which mediates facial mechanosensation and nociception^4,9^. More posteriorly, the otic placode invaginates to form the otic vesicle, giving rise to the inner ear labyrinth and the vestibulocochlear ganglion^10,11^. The epibranchial placodes - geniculate, petrosal, and nodose - produce visceral sensory neurons of the cranial ganglia associated with the facial (VII), glossopharyngeal (IX), and vagus (X) nerves, respectively^12^. Collectively, these placodes generate much of the vertebrate cranial sensory apparatus, underlying vision, hearing, olfaction, equilibrium, somatosensory and viscerosensory perception^13^.

Cranial placodes arise from the neural plate border (NPB), a transient region of the early embryonic ectoderm located between the neural plate and the non-neural epidermis^8,14–16^. During neurulation, the NPB is patterned by intersecting gradients of signaling molecules such as BMP, FGF, and WNT, which together subdivide the ectoderm into neural, neural crest, pre-placodal, and epidermal territories^17^. Within the NPB, the pre-placodal region (PPR) emerges as a specialized ectodermal domain characterized by the expression of conserved transcription factors including Foxi3^18,19^. Subsequently, the PPR becomes regionalized along the anterior-posterior axis to generate the distinct cranial placodes that will form the sensory organs and ganglia of the head^13,20^.

The cellular and molecular logic by which naïve ectodermal cells at the neural plate border acquire pre-placodal identity remains obscure^20^: it is unclear whether placodal specification follows a deterministic sequence of lineage bifurcations or proceeds through probabilistic fate bias within a multipotent ectodermal field. Competing models - including the neural plate border model, binary competence model, and probabilistic model - offer different interpretations of how NPB cells resolve toward placodal, neural crest, or epidermal fates^17,20–22^. The transcriptional and signaling codes that govern this diversification are only partially defined: although Foxi, Six, Eya, Pax, and Dlx families have been implicated^13,19,23^, the complete gene regulatory network and intercellular signaling mechanisms dictating individual placodal identities remain poorly understood. Moreover, it is unclear how discrete placodal territories are established within the continuous ectodermal sheet of the PPR, resulting in distinct placodes with well-defined boundaries. Do emerging placodes arise by competitive recruitment of progenitors from a common pool, or by independent local specification events?

Questions surrounding the specification of cranial placodes naturally extend into broader evolutionary questions about how the diverse placodal types originated in early vertebrates. If, during development, the pre-placodal ectoderm behaves as a transcriptionally graded field in which adjacent placodes gradually segregate from multipotent progenitors, this organization may recapitulate an ancestral condition in which all placodal derivatives evolved from a single, broadly sensory-competent ectodermal domain. Conversely, if each placode exhibited an early and transcriptionally discrete identity governed by specific combinations of transcription factors and signaling pathways, it would suggest that individual placodes evolved as semi-independent modules, perhaps through repeated co-option of the same pre-placodal gene regulatory toolkit in different ectodermal contexts. Our incomplete understanding of the transcriptional and signaling code that defines individual placodal identities therefore has implications for the evolutionary question of how and when these molecular modules diversified in the vertebrate lineage. To answer these long-standing questions, we constructed a composite single-cell, spatial, and clonal atlas of the mouse cranial placodes, capturing their molecular and lineage architecture at cellular resolution (see the schemes of the approach in Fig. 1). This approach enabled us to obtain new insights into placodal specification and revisit classical evolutionary models of placode origin through the lens of developmental dynamics and cell-state transitions in the mammalian embryo.

**Figure 1.**
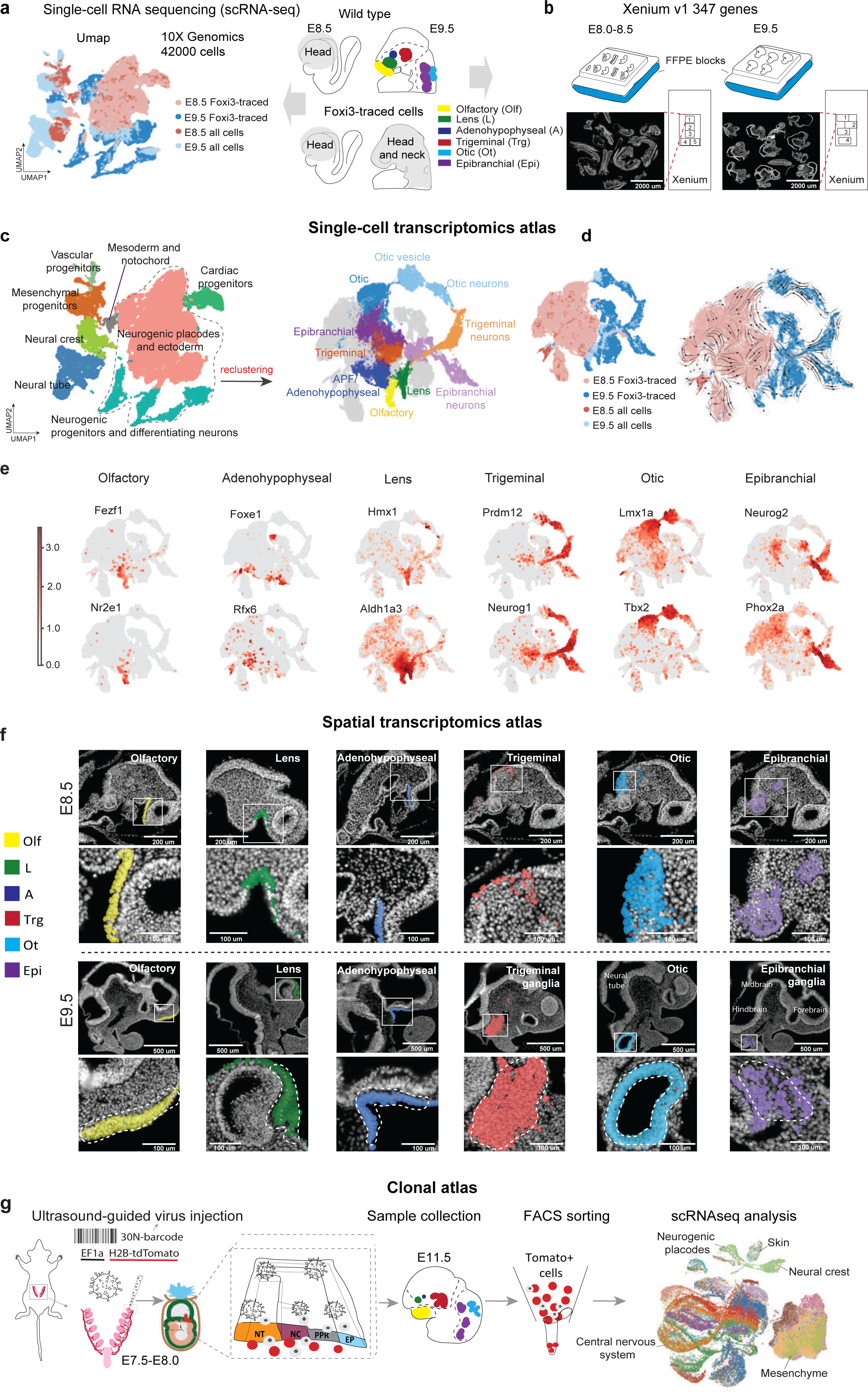
Atlases of neural plate border derivatives including neural crest and cranial placodes. **(a)** The concept of a single cell transcriptomics atlas: left, UMAP embedding of sequenced cells. Right, schematic of the experimental procedure: dissection of wild type (top) and Foxi3-traced (bottom) cranial placodes at E8.5 and E9.5. **(b)** The concept of a spatial transcriptomics atlas: strategy for sectioning E8.5 and E9.5 embryos prior to analysis by Xenium v1 347 genes. **(c)** Left, UMAP embedding showing annotation for major clusters including neural plate border derivatives. Right, UMAP embedding contains only epithelial-like (EPCAM+) cells leading to identification of pre-placodal and placode-containing clusters. **(d)** Analysis of RNA Velocity reveals dominant paths of cellular state transitions in transcriptional space, with arrows indicating progression from present transcriptomic states to predicted future states. **(e)** UMAP plots for key gene markers known to be the features of different cranial placodes. **(f)** Spatial representation of clusters containing cranial placodes detected by the Xenium platform, for the E8.5 (top) and E9.5 (bottom) staged embryos. **(g)** The concept of a clonal atlas: experimental workflow for massive parallel clonal labelling and tracing experiment using NEPTUNE high frequency ultrasound and microinjection system. Embryos were transduced *in utero* with barcoded lentivirus at E7.5-8. The traced embryos were harvested at E11.5 for cell dissociation and single cell library preparation.

## Results

### Single cell transcriptomics and spatial atlases of the cranial placodes and neural plate border region

To understand the development of cranial placodes and resolve the deterministic vs probabilistic hypothesis of placode formation, we built an integrated atlas of single cell transcriptional states, cell positional behavior, and progenitor multipotency. We used a combined approach: single cell RNA-seq of dissociated cells and spatial transcriptomics by Xenium v1, which together help discern gene regulation and spatial interactions, and *in vivo* single-cell barcode labeling for clonal analysis to reveal lineage histories, hierarchies and segregation.

First, to define the transcriptional dynamics and gene regulatory networks underlying placode formation, we performed single-cell transcriptomic analysis of wild-type mouse cranial development spanning neural and non-neural ectoderm and their major derivatives, including neural tube, neural crest, placode-derived sensory lineages, and emerging embryonic epithelia. To enrich this atlas for placodes (a rare cell population), we took advantage of Foxi3-CreER mice, which recombine in early embryonic ectoderm of the neural plate border, and predominantly label cranial placodes and ectoderm when tamoxifen is administered at embryonic days (E) 6.5 and E7.5^23,24^. The cells from wild-type and Foxi3-Cre traced embryos were co-clustered together to show developmental processes at E8.5 to E9.5 (Fig. 1a-c). To isolate and better understand the development of cranial placodes, we isolated all cells expressing *Epcam*^25,26^, as well as their immediate derivatives (neuronal differentiation trajectories), and re-clustered the resulting set of cells to increase clustering resolution and better visualize placodes and possible transitions (Fig. 1c, Supplementary Fig. 1). RNA Velocity analysis validated our basic interpretations of how the placodally-derived progeny differentiate – particularly in the otic vesicle and cranial ganglia resulting from differentiation trajectories originating in corresponding placodal fields (Fig. 1d). Placodal cells clustered together and expressed known markers: for example, at E8.0-8.5, placodal cells expressed canonical pre-placodal ectoderm markers such as Six1 and Eya1. The combined dataset resolved established markers for all six cranial placodes: olfactory (*Foxg1, Dmrt3, Nr2e1, Fezf1, Sp8*), lens (*Hmx1, Pcdh9, Aldh1a3, Mgarp, Thrb, Six3, Six6, Pax6*), adenohypophyseal (*Pitx1, Pitx2, Lhx3, Foxe1*), trigeminal (*Pax3, Wnt6, Tfap2b, Neurog1*), otic (*Oc90, Pax2, Sox10, Lmx1a, Tbx2*), and epibranchial (*Pax2, Pax8, Shisa6, Nrp1, Fgf4, Fgf3, Fgf15*) (Fig. 1e). The olfactory and adenohypophyseal placodes were not fully segregated in our clustering analysis especially at day 8.5, as intermediate cell states were observed. We therefore refer to this region as the anterior placodal field (APF)/adenohypophyseal, as this cluster is enriched in adenohypophyseal placode cells with a smaller contribution from the olfactory placode. Similarly, the olfactory cluster likely includes some cells that will end up in the adenohypophyseal placode.

At E9.5, and already during the transition from E8.5 to E9.5, we observed sharper molecular segregation of placodal territories and their derivatives, with more clearly defined olfactory, lens and otic vesicle populations, as well as ongoing neurogenesis arising from trigeminal, otic and epibranchial placodes (Fig. 1c-e). The other annotated clusters correspond to a range of non-neural epithelial subpopulations, including branchial arch ectoderm, pharyngeal endoderm, and cardiac-associated epithelia.

To understand the spatial context of placodal development, we next took advantage of Xenium v1 spatial transcriptomics (347-gene hybridization panel), which helped to locate gene expression signatures observed in the single cell atlas at both E8.5 and E9.5 stages (Fig. 1f). Finally, to understand the cell fates arising from progenitors at the borders of developing ectodermal structures, we generated a clonal atlas based on massively parallel DNA/RNA barcoding via *in utero* microinjections of lentiviral particles (Fig. 1g).

Previously, the characterization of cell dynamics at the developing neural plate border region in chick embryos revealed that progenitors are in a mixed cell state and likely resolve the neural tube, neural crest, placodal and epithelial fates probabilistically over some period of time^26^. Similarly, in mouse embryos we observe the main lineages separating over 24-48 hours. At E8.5, the neural tube fates are closely connected to the neural crest, whereas the placodes are embedded into a continuum of epithelial cell states in which the non-neural ectoderm dominates, in agreement with chick data^25,26^, (Fig 2a, Supplementary Fig. 2a). The faint bridge of intermediate cells connecting only the anterior placodes (*Vax1/Fezf1*) + epithelium with the neural tube + neural crest is present as early as E8.5 (Fig. 2b, Supplementary Fig. 2b-d). This suggests that in mouse embryos, the first lineage decision splits the neural tube ectoderm and the placodes, whereas generation of the neural crest cells and their subsequent separation from the neural tube happens much later in development. However, placode cells are not clearly transcriptionally separated from each other at the borders of UMAP clusters or spatial populations, nor from the non-neural ectoderm, prior to the production of placodal derivatives (summarized in Fig. 2c).

**Figure 2.**
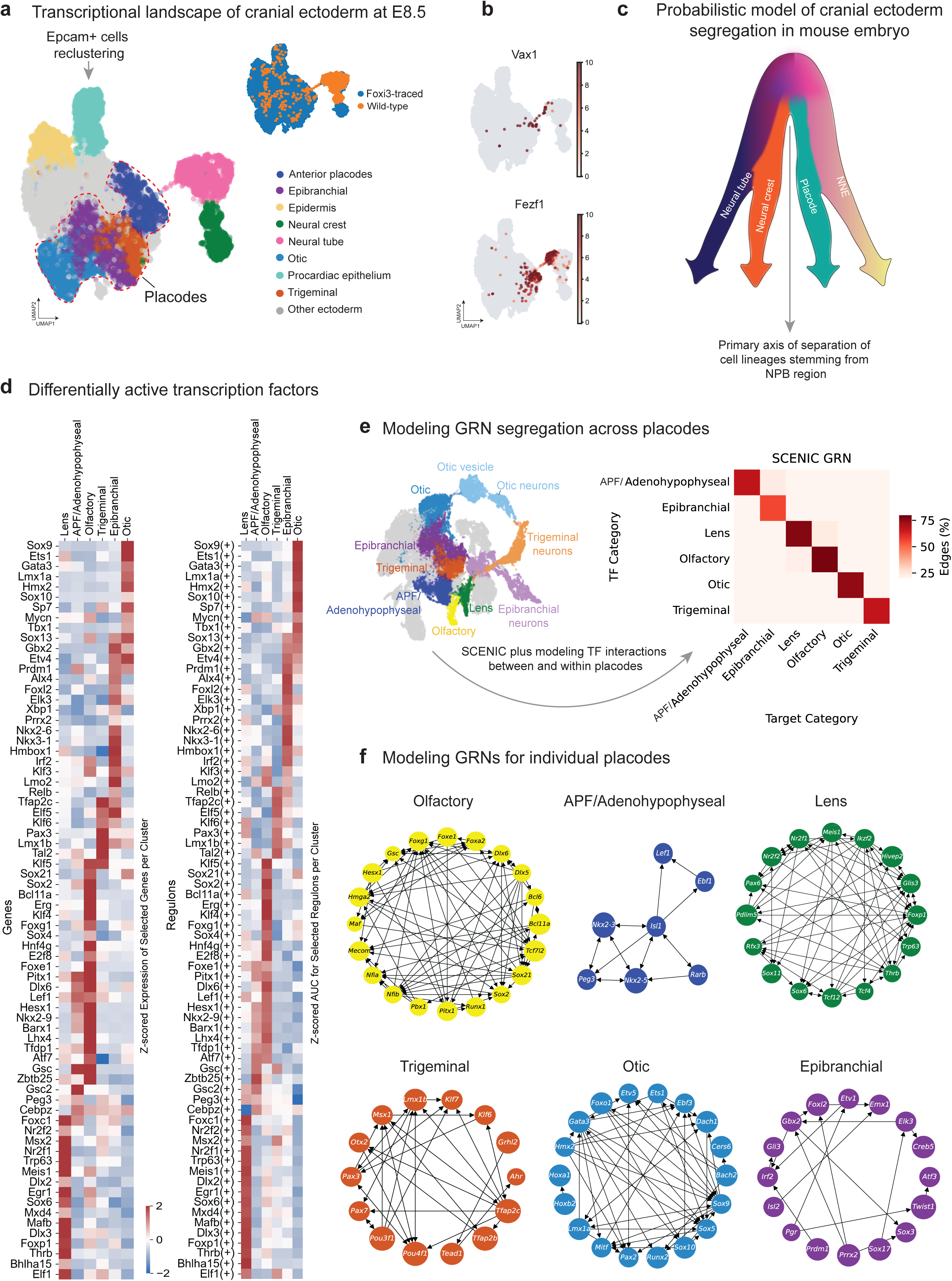
Segregation of lineages originating from the neural plate border. **(a)** UMAP visualization of merged *Foxi3*-traced and wild-type datasets at E8.5. Only *Epcam+*, neural crest and neural tube populations are selected for this clustering and UMAP generation. **(b)** UMAP feature plots for *Vax1* and *Fezf1* showing expression in the anterior preplacodal area, bridge, and the neural tube. **(c)** Schematics approximating the hierarchy of lineage segregation in the neural plate border region in mouse and chick embryos. **(d)** Heatmap showing the expression of placodal cluster-specific transcription factors and active regulons detected via SCENIC. **(e)** Left, UMAP embedding showing cluster annotation. Right, heatmap showing the degree of transcriptional cross-interactions (via transcriptional activation) between individual placodal cluster GRNs. **(e)** Reconstructed and modelled placodal cluster GRNs form discrete subnetworks at E8.5. The model infers transcription factor interactions within individual GRNs.

Next, we derived the sets of transcription factors (TFs) and their regulons (gene modules coordinated by a single TF) that characterize each placode (Fig. 2d, Supplementary Fig. 3a-f). Because we observed distinct TF expression patterns across individual placodes, we next attempted to reconstruct the GRNs governing the segregation of these distinct fates. Using SCENIC, we inferred separate GRNs for the early and late placodal cells in our atlas at E8.5 and E9.5, respectively. Mirroring the observed TF expression patterns, the reconstructed GRNs displayed strong segregation between the subnetworks for each placode (Fig. 2e, f). Specifically, the TFs that were highly expressed in the cells from a specific placode were predicted to primarily co-regulate other TFs that were also highly expressed in the same cells. However, this inference cannot capture cross-repression events, which likely underlie border refinement and competitive fate allocation where different epithelial domains are in contact with each other. We also took advantage of the STRING^27^, which highlights interacting, co-operating, and co-occurring proteins using previously published data. In this analysis, we generated smaller connected networks of interacting TFs in every placode (Supplementary Fig. 3g). This suggests that such interacting TFs might play important roles in the placode-specific GRNs.

Although we primarily focused on reconstructing and modelling GRNs for individual placodes prior to formation of derived cell types, we also analyzed the neurogenic trajectories leading to neuronal subtypes arising from the trigeminal (Supplementary Fig. 4), otic (Supplementary Fig. 5), and epibranchial placodes (Supplementary Fig. 6). As expected, the well-known and ganglia-specific neurogenic transcriptional factors appeared to be transcriptionally active at early stages of neurogenesis according to the analysis of regulons (via SCENIC) (Supplementary Fig. 4c-5c-6c). These maps are useful to predict possible signaling pathways (for example, Wnt signaling mediated by *Lef1* in the trigeminal placode) and other factors driving the specificity of neuronal differentiation in each placode. For instance, we observed differentially active regulons corresponding to positional anterio-posterior codes, such as Dlx3(+), Tbx3(+), Hoxc4(+) and others (Supplementary Fig. 4c-5c-6c).

Next, we attempted to map the possible molecular interactions in placodes, which are essential to decipher the regulatory mechanisms driving early development of placodal identities. Our analysis shows that every placode produces a set of specifically expressed ligands, for which the cognate receptors are present in other placodes and non-placodal adjacent structures (Supplementary Fig. 7 and 8, Supplementary Table 1). Examples of specifically expressed ligands include the following - for olfactory placode: *Dlk1, Cxcl12, Fgf16, Wnt5b*; lens: *Sema3a, Bmp4, Bmp7*, *Fn1*; APF/adenohypopheseal: *Ltbp1, Ybx1, Nrxn1*; trigeminal: *Alcam, Bmp2, Bmp5*; otic: *Ptn, Tgfa*; epibranchial: *Fgf4, Jag1, Has2.* The specifically expressed receptors include - olfactory: *Lgr4, Fgfrl1, Fzd5*; lens: *Cnr1, Antxr1, Met*; APF/adenohypopheseal: *Rack1, Nrxn3, Ptprg*; trigeminal: *Sdc4, Ror2, Erbb2*; otic: *Bmpr1rb, Epha4, Robo1*; epibranchial: *Fzd7, Nrp1, Fgfr1 (*Supplementary Fig. 8). Of note, although ligand-receptor interactions between placodes in the plane of the epithelium may refine individual placodes, additional signals come from outside the ectoderm (e.g. mesoderm or neural plate). The validation and refinement of the predicted interactions (Supplementary Table 1) is a goal of future experimental efforts.

Despite not being fully separated in the UMAP, different placodes occupy various distinct positions in the UMAP which resemble the anterior-posterior axis of the embryo, with more anterior placodes such as olfactory, lens and adenohypophyseal appearing in the UMAP next to each other (Fig. 1 and 2). This analysis of the UMAP and clustering allowed us to hypothesize that placodes do not emerge as fully-separated entities in epithelial space but rather form from gradual border refinement starting from intermediate transcriptional states visible at E8 (posterior placodes) and E9 (anterior placodes) (Fig. 3a). This is supported by the comparative analysis of expressed transcription factors showing more indistinct and party overlapping profiles and their corresponding regulons (Fig. 3b): the expression of transcription factors covers a domain with gradually fading borders, which overlap with similarly fading borders of the neighboring placodes and other epithelial structures. However, the transcriptional activity of these transcription factors (regulons) appeared to be more confined towards placodal centroids and largely non-overlapping with regulons driving the development of neighboring placodes (depicted in Fig. 3b in the scheme on the right). This implies that placodes segregate from each other through competitive interactions of patterning transcription factors.

**Figure 3.**
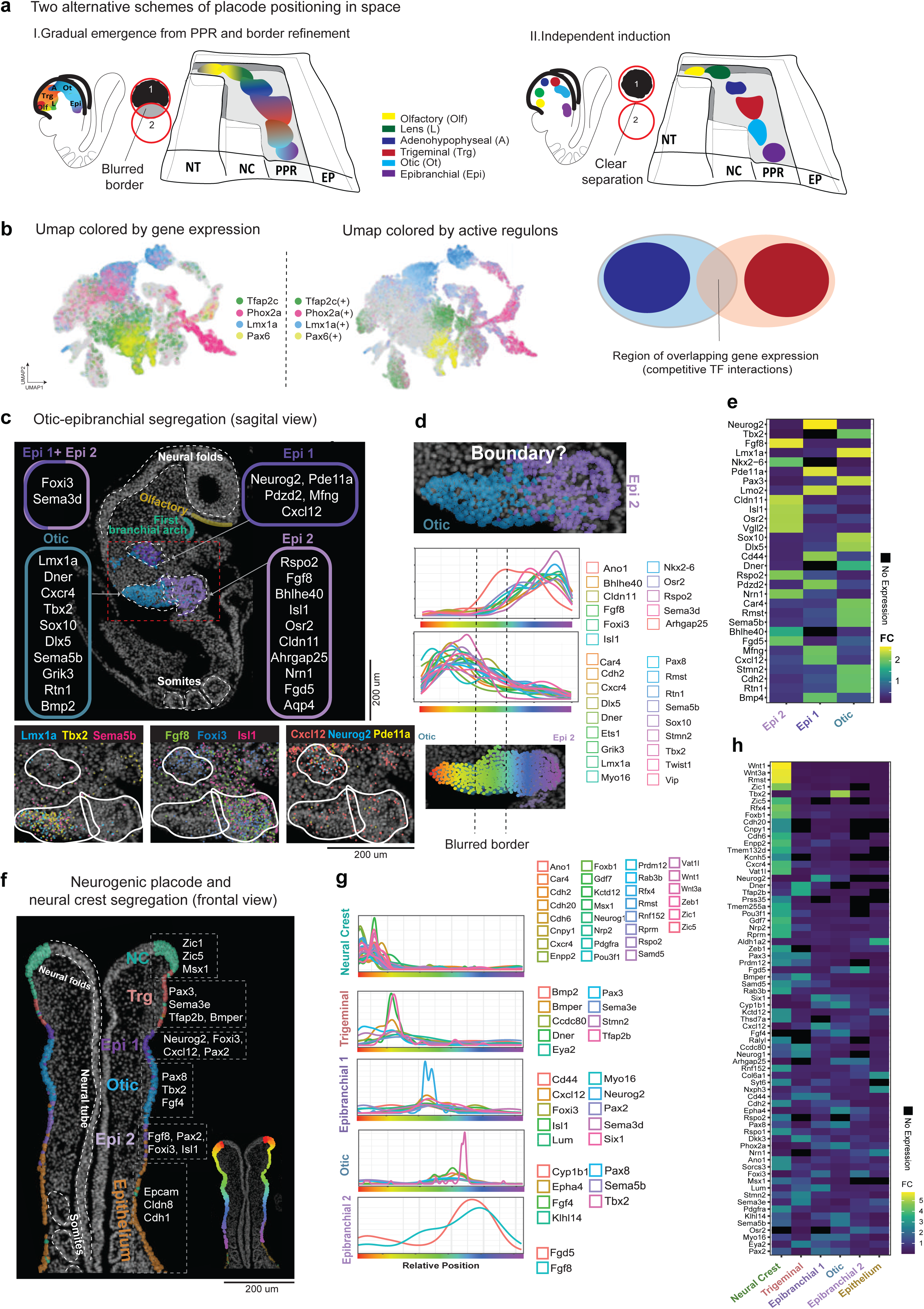
Fuzzy boundaries and continuity of epithelial cell states revealed by single-cell and spatial transcriptomics. **(a)** Two alternative schemes of placode initiation within the ectodermal space. Left, probabilistic and competitive induction along the axial gradient – with fuzzy boundaries, which undergo gradual refinement. Right, isolated induction model – all placodes are spaced out since the beginning. **(b)** UMAP plots of gene expression of key transcription factors and their active regulons for Tfap2c, Phox2a, Lmx1a, and Pax6, illustrating the differences between broad and fuzzy transcription factor expression and more compacted regulon activity. **(c)** Spatial segregation of otic and epibranchial placode-including clusters and corresponding expressed genes in E8.5 mouse embryo. The stars next to gene names show the rating of specificity of a selected marker. **(d)** Fuzzy boundary between otic and epibranchial placode-containing clusters analyzed at the level of differentially expressed marker genes. Line graphs show individual gene expression levels of otic- and epibranchial- cluster genes over two interlaced placodal regions. **(e)** Heatmap showcasing thirty key variable genes characterizing the three placodal regions – otic, epibrachial 1 and epibranchial 2. **(f)** Spatial atlas shows the position and markers of all major epithelial and epithelia-derived cell states at E8.5: the neural crest, trigeminal, epibranchial, otic and epithelial regions. The inset shows how color code in the spatial data corresponds to the color legend for line graphs in g. **(g)** Line graphs and the heatmap **(h)** showing the characteristic genes for cell populations in **f** over the linearized epithelium.

We took advantage of our spatial transcriptomics atlas to clearly demonstrate the existence of blurred boundaries and spatial overlaps between neighboring placodes (Fig. 3, Supplementary Fig. 9-12). Xenium spatial transcriptomics enabled single-cell resolution and segmentation, allowing accurate localization of transcriptional programs at placodal borders (Fig. 3c, f). We used the Xenium mouse brain panel supplemented with a custom set of additional 100 genes (see Supplementary Table 2), which were selected based on our single cell transcriptomics sequencing and canonical markers of placodes ^28–30^. We identified spatially restricted transcriptional programs encompassing the entire cranial ectoderm, which allowed us to visualize all territories including neural plate/tube, neural crest, placodal epithelium, and epidermis. This confirmed that these domains are molecularly distinct by mid-neurulation (Supplementary Fig. 10-12). Next, we generated UMAP embeddings from the Xenium data to better visualize and annotate all placodal and other epithelial clusters (Supplementary Fig. 10-12). As before, we selected *Epcam*-positive cells and re-clustered them to capture additional cellular diversity within the epithelial population at E8.5, independently analyzing three sections of a single paraffin block with 13 embryos embedded in different orientations (Supplementary Fig. 10-12). The analysis of segmented cells and their transcriptomes confirmed the presence of placodes and neighboring structures, consistent with our single cell transcriptomics performed on dissociated cells (Fig. 2).

The unbiased placodal territories that we demarcated as the result of finding conventional markers being expressed in specific unbiased clusters turned out to be relatively large and in contact with each other, according to anterior-posterior and dorso-ventral coordinate logic^31,32^, (Fig. 3, Supplementary Fig. 9-12). As a result, in addition to positioning the individual placodes, we were able to capture mixing transcriptional programs, where pairs of placodes meet in 2D-epithelial space (Fig. 3c-e). The contact points between other placodes also revealed gradual gene expression transitions within the placodal epithelium (Fig. 3f-h, Supplementary Fig. 9-12). This suggests the existence of mixed-identity populations at the forming boundaries at E8.5, i.e. cells that have characteristics of neighboring placodal or other territories. These “boundary cells” might represent either bipotential or multipotent progenitors that have not yet resolved which placode or other epithelial structure to join (Fig. 3a-b). Alternatively, this population might later take on a non-neural epithelial fate expanding the domain of the future skin. Indeed, our data show that this blurred-boundary logic can be detected in placode-epidermis/epithelium border territories (Supplementary Fig. 9-12). However, when we analyzed the spatial transcriptomics data from E9.5 embryos - when placodal derivatives are already emerging (4 sections with multiple embryos embedded within a single paraffin block), we detected clean and well-separated borders between posterior placodes (trigeminal, otic, epibranchial), as well as between neural tube, cranial neural crest and cranial epidermis (Supplementary Fig. 13). At E9.5, the anterior placodes remain in close contact with each other, sharing blurred transcriptional programs at borders where the olfactory placode meets either presumptive lens or adenohypophyseal epithelium^33^, (Supplementary Fig. 13).

### Clonal atlas of the cranial placodes and other related epithelial structures

To construct a comprehensive clonal atlas of the placodes, surface ectoderm, neural tube and neural crest, we combined ultrasound-guided *in utero* delivery of barcoded viral libraries with high-resolution lineage reconstruction – an approach recently taken to resolve the clonal logic of cochlear development^34^, and neural crest specification^35^. Using the NEPTUNE platform^36^, we performed microinjections of a TREX lentiviral barcode library directly into the amniotic cavity of E7.5-E8.0 mouse embryos (Fig. 1g). The TREX library comprises millions of uniquely barcoded viral constructs^35,37,38^. Upon transduction, ectodermal progenitors stably integrate unique barcodes, which are inherited by each daughter cell. By sequencing these barcodes from head and neck single-cell transcriptomes collected at E11.5, we reconstructed clonal genealogies spanning neural crest, neurogenic placodal, neural ectodermal, and non-neural ectodermal lineages. We recovered 171,093 sequenced cells with clonal barcodes, capturing the major embryonic compartments that contribute to cranial structures (Fig 4a-c). In the resultant embedding, cell clusters were observed corresponding to the central nervous system (CNS), mesenchyme, neural crest, and an ectodermal domain containing cranial placodes and developing epidermis (skin) (Fig. 4a, Supplementary Fig. 14).

**Figure 4.**
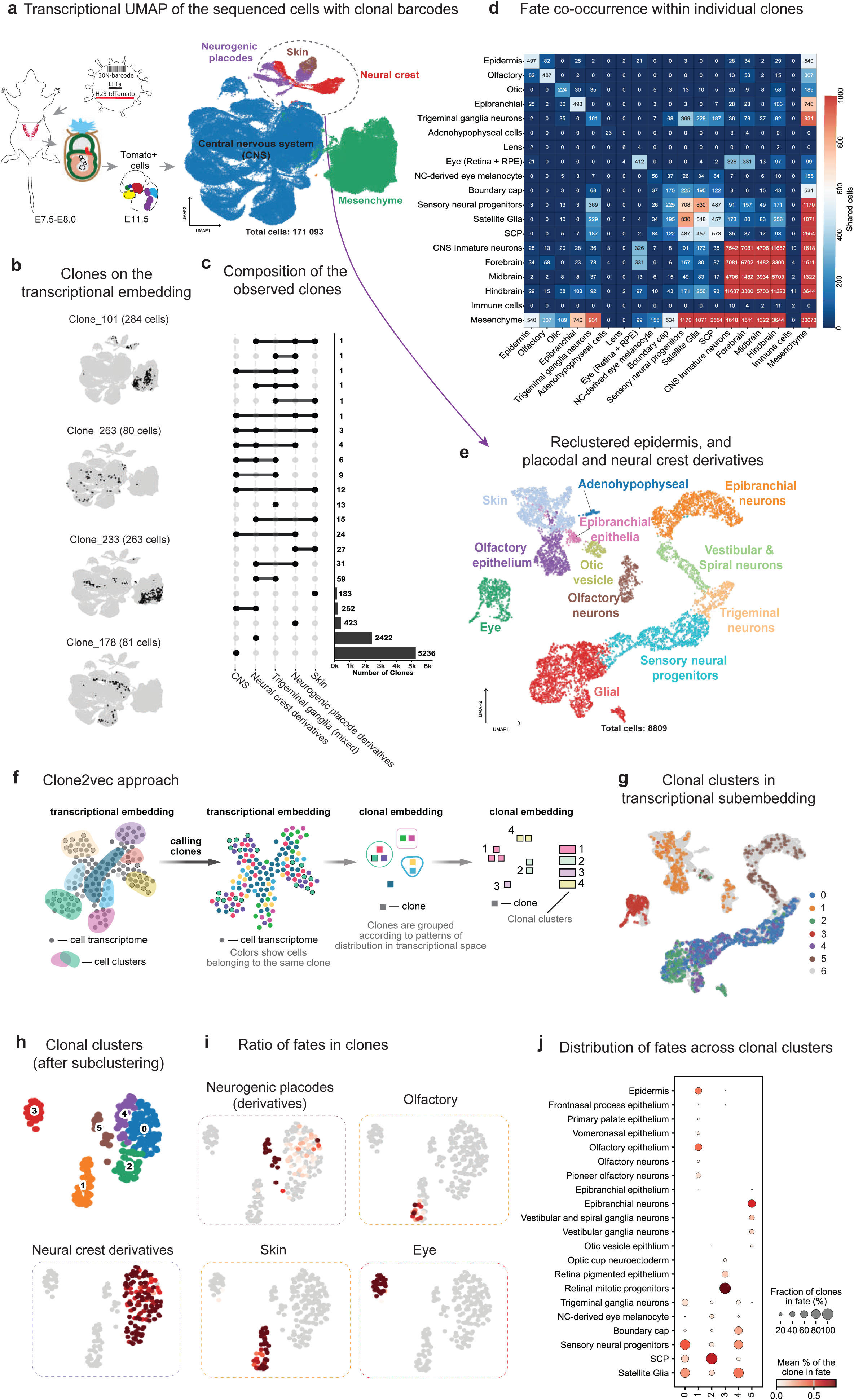
Clonal atlas of derivatives of the neural plate border region with focus on cranial placodes. **(a)** UMAP embedding of the integrated clonally-barcoded dataset (resulting from injections of TREX lentiviral library at E7.5-8) generated from traced E11.5 mouse embryos. Note major clusters derived from the neural crest, neuronal, non-neuronal and pre-placodal regions. **(b)** Examples of individual clones identified and shown in the transcriptional embedding. **(c)** Upset plot showing the principal composition of the observed clones. **(d)** Heatmap showing the fate co-occurrence matrix based on shared clones. **(e)** UMAP of reclustered placodal and neural crest neuro-glial derivatives. **(f)** Schematic illustration of Clone2Vec approach. **(g)** Mapping of all observed epithelial/neural crest-derived neuroglial clonal clusters on the corresponding transcriptional embedding. **(h, i)** Clone2vec clonal embedding of the same subset of cells, represented as a UMAP of clones, and color coded by the major clonal clusters observed. **(j)** Dot plot shows the distribution of fates across clonal clusters of derivatives of cranial placodes, neural crest, and embryonic epidermis.

Individual clones showed diverse distribution patterns: some were tightly localized within a single region in the UMAP transcriptional embedding, whereas others dispersed across adjacent territories or related derivatives, consistent with lineage progression through intermediate states (Fig. 4b). Summarizing clone compositions across broad developmental domains revealed that most clones were fate-restricted at the level of major compartments (e.g., CNS-only, neural crest-only, placode-only, or ectoderm/skin-only), while a smaller subset spanned multiple related compartments, suggesting early common progenitors (Fig. 4c). Quantitative analysis of clonal composition showed that CNS-related clones were the most abundant (5,236 clones), followed by the cranial neural crest derivatives (2,422 clones). Placodal clones (423) and epidermal clones (183) were less abundant, which is expected, given the relative sizes of neural plate versus non-neural ectoderm in the anterior region of a mouse embryo at E7.5.

We next quantified lineage coupling between annotated fates using fate co-occurrence and clonal co-occurrence matrixes based on shared clonal membership. This analysis consistently confirmed strong, though not absolute, within-domain fate connections within clones, for example, among CNS subdivisions (forebrain, midbrain, hindbrain, immature neurons) or neural crest derivatives (mesenchyme etc.), all consistent with known shared ancestry within the neural plate and neural plate border territory (Fig. 4d, Supplementary Fig. 15). Beyond these canonical relationships, the co-occurrence structure also revealed a subset of clones spanning major ectodermal domains. Importantly, clones contained both CNS and neural crest derivatives more often as compared to placodal derivatives, which supports the closer connection between the neural crest and developing neural tube as compared to the neural tube or neural crest and the placodes (Fig 4c-d, Supplementary Fig. 15). The placodes, in contrast, contained more clones shared with the developing neural crest, the epidermis (future skin), and with each other. Notably, we observed substantial clonal sharing between the olfactory placode and non-neural epidermis. This pattern points to the existence of incompletely specified epithelial progenitors around the time of lentiviral infection (E7.5-E8.0) that were at least bipotent, contributing both to olfactory lineages and to facial skin. In contrast, clones linking the olfactory placode to anterior CNS fates were present but less frequent, suggesting either a narrower window for shared ancestry or more rapid lineage segregation at the anterior neural/placodal interface. We also detected frequent co-occurrence among posterior placodal derivatives - particularly trigeminal, otic, and epibranchial lineages - consistent with the hypothesis suggesting the existence of neighboring multipotent or undecided progenitor states at the time of virus administration. Unfortunately, beyond the olfactory placode, the other anterior placodal lineages were far less represented in our clonal atlas: lens and adenohypophyseal derivatives were insufficiently labeled, likely reflecting limited number of their progenitors at the labeling stage. Together, these support a model of gradually refined blurred borders presented in Fig.3a-left. Consistent with this, despite the observed “preferential” clonal connections constrained within the major lineage domains, some individual clones contained both placodal derivatives and the neural crest derivatives, with a fraction of clones including CNS-resident cells, indicating that some barcoded progenitors were multipotent and had not yet resolved into strict domain-specific lineages at the time of labeling (at E7.5-E8.0). A good example includes the olfactory placode, which shares many clones with epidermis and some clones with the forebrain cell types (Fig. 4c-d and Supplementary Fig. 15).

To reduce the complexity and improve the resolution of clonally labeled placodal derivatives and related cell populations, we re-clustered the relevant parts of embedding in Fig. 4a. The new refined embedding better resolved placode-derived fates at E11.5, including olfactory epithelium and olfactory neurons, adenohypophyseal cells, epibranchial derivatives including diverse neurons, the otic vesicle and related cell types such as vestibular/spiral ganglion neurons, trigeminal ganglion neurons, other sensory neural progenitors stemming from the neural crest (anterior DRGs), melanocytes, peripheral glial populations including Schwann cell precursors and boundary cap cells, alongside neighboring skin and eye-associated cells (utilized as a negative unrelated lineage control) (Fig. 4e, Supplementary Fig. 16-17). Furthermore, to extract the stereotypical “clonal programs” and fate biases in placodal and neural crest derivatives, we implemented our new Clone2Vec strategy^35^, that represents each clone by its distribution in transcriptional space and embeds clones according to similarity of those distributions (explained in Fig. 4f). More generally, Clone2Vec learns clonal context by identifying pairs of clones that share nearest-neighbor cells in the transcriptional landscape. As a result, Clone2Vec reveals the most stereotypical aspects of clonal behavior aggregating them into the most reliable patterns. The resulting clonal embedding identified six discrete clonal clusters (0-5, 6 is for negative cells) (Fig. 4e) whose members mapped to coherent regions of the transcriptional sub-embedding in Fig. 4g, implying that clones mainly fall into recurrent lineage-pattern classes (of course, these major classes result from the most stable and re-occurring clonal structures and do not reflect less recurrent or rare clonal situations).

Consistently, clones contributing to cranial placodes, olfactory, neural crest derivatives, skin, or eye localized to the corresponding transcriptional territories, providing a direct visualization of fate-biased clone structure within the atlas (Fig. 4h-i). Finally, the deeper analysis of more granular fate composition of each clonal cluster further demonstrated that clonal clusters form as a consequence of biased lineage outputs: some clusters were biased for specific placode derivatives (for instance, olfactory neurons) and skin - clonal cluster 1, whereas the others showed the distribution of neural crest-derived cell types (Fig. 4j, Supplementary Fig. 15). Importantly, the close connectedness of the olfactory placode and developing skin – for example, in clonal cluster 1 in Fig. 4j suggests the existence of a pool of common progenitors specifically in the anterior part of an embryo. At the same time, clonal clustering did not reveal any significant connection of the olfactory placode and the derivatives of the posterior placodes - as expected due to the spatial positioning logic. Another key result of this analysis is that the posterior placodes appeared clonally interconnected with each other – being organized into the clonal cluster 5 in Fig. 4j. Together, these results appear consistent with the ideas of partly separated anterior and posterior pre-placodal territories.

### Implications for the evolutionary origin of placodes

Mapping gene regulatory networks and transcriptional profiles across placodes enables the identification of shared regulatory cores and placode-specific differences. These comparisons can uncover ancient transcriptional networks while clarifying the mechanisms that drove placodal diversification, providing a framework for understanding both their evolutionary origin and specialization. To investigate the comparative regulatory logic of cranial placodes, we first quantified pairwise overlap of expressed genes across six mouse placodes (adenohypophyseal-APF, epibranchial, otic, olfactory, lens, trigeminal). This analysis uncovered extensive sharing of genes among the individual placodes (Fig. 5a). These genes were initially identified as the differentially expressed set of placodal genes obtained in comparison of all placodal clusters together vs other clusters. The largest overlap was observed between olfactory and lens placodes (1410 shared genes), indicating unusually strong similarity in protein usage between these two placodes. Substantial overlap was also apparent among other placode pairs (e.g., olfactory-otic, otic-lens, trigeminal-lens; Fig. 5a), whereas a smaller, but still prominent, set of shared expressed genes connected all placodes, supporting the idea of a shared “placodal” regulatory network with placode-specific elaborations.

**Figure 5.**
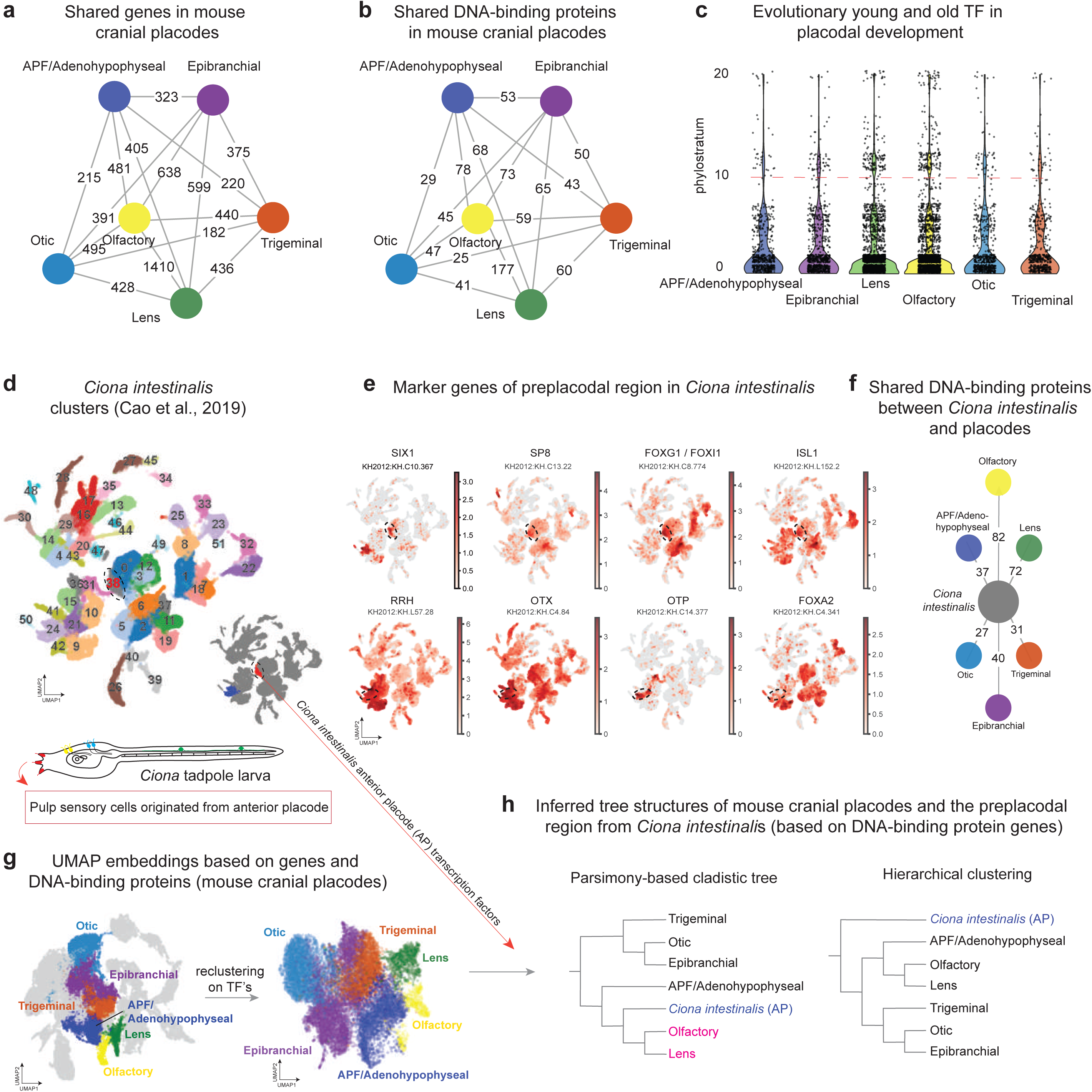
Comparative single cell transcriptomics analysis of tunicate and mouse placodes. **(a)** Visualization of shared genes and DNA-binding proteins **(b)** expressed in mouse cranial placodes. **(c)** Phylostratigraphic analysis: evolutionary young and old transcription factors in placodal development. **(d)** UMAP visualization of single cell data previously derived from developing *Ciona intestinalis* larva (Cao et al., 2019). **(e)** UMAP plots of expressed marker genes labelling proto-placodal regions (anterior –cluster 38 and more posterior – cluster 24) in larval *Ciona intestinalis*. The anterior proto-placodal region giving rise to palp sensory cells is highlighted by the dotted line in UMAPs for Six1, Sp8, Fox, and Isl1 genes. **(f)** Visualization of numbers of shared DNA-binding proteins between anterior proto-placodal area of larval *Ciona intestinalis* and mouse cranial placodes. **(g)** Reclustering of the mouse neural plate border derivatives dataset (left) based exclusively on the expression of DNA-binding proteins (right). **(h)** Left: Cassiopeia-built parsimony-based tree derived from a set of DNA-binding proteins expressed in *Ciona* anterior placode in relation to mouse cranial placodes. Right: tree based on hierarchical clustering of cluster-specific transcriptional states. AP, anterior placode; APF, anterior placodal field; TF, transcription factors.

We next asked whether this degree of sharing reflects broad transcriptional similarities or reflects a stereotyped use of few DNA-binding proteins to direct similar regulatory mechanisms. When we repeated the overlap analysis using only the shared DNA-binding protein set (rather than using all differentially expressed genes), as expected, pairwise intersections were markedly smaller (maximum overlap 177 DNA-binding proteins between olfactory and lens; Fig. 5b). Most other placode pairs shared tens of such regulatory genes (Fig. 5b). This implies that placode identity may rely less on unique transcription factors and more on combinatorial reuse^39,40^, context-specific cofactor deployment, and downstream target selection.

Because conserved regulatory networks often coexist with lineage-specific innovations, we analyzed the evolutionary age distribution of transcription factors involved in placodal development using phylostratigraphy. This approach^41^, assigns genes to evolutionary age classes, allowing us to infer when they emerged and when they were likely recruited into placodal identity programs. Across all placodes, transcription factorsets were dominated by evolutionarily old transcription factors, but with long upper tails corresponding to younger, lineage-restricted transcription factors (Fig. 5c). This pattern suggests that individual placodal programs are built on an ancient, conserved regulatory core supplemented by a limited set of younger components. Such a configuration is consistent with progressive remodeling of a shared developmental framework rather than the de novo invention of entirely new gene modules in each placode. Notably, the construction of placodal gene regulatory networks involves extensive recruitment of ancient genes alongside younger additions, underscoring the predominance of evolutionary reuse over novelty.

To test if elements of the placodal regulatory network predate vertebrates, we leveraged published single-cell transcriptomic data from the ascidian *Ciona intestinalis* (Fig. 5d) and identified a candidate “anterior proto-placodal region” by marker gene expression using previously published studies^42–44^. Within the *Ciona* atlas, this discrete transcriptional domain expressed canonical preplacodal/placodal regulators, including SIX1, SP8, FOXG1/FOX11, and ISL1, together with additional markers (RRH, OTX, OTP, FOXA2) that in combination delineate this region within the broader embryo-wide embedding (Fig. 5e). These marker patterns support the existence of an anterior *Ciona* cell population with a placode-like transcriptional signature, providing an entry point for cross-species comparison of regulatory factor usage.

We then analyzed the overlap between the sets of DNA-binding proteins expressed in the *Ciona* proto-placodal-like region and each mouse placode (Fig. 5f). The strongest overlap was observed with olfactory (82 shared DNA-binding proteins) and lens (72), followed by epibranchial (40), adenohypophyseal (37), trigeminal (31), and otic (27).

Consistent with this result, dimensionality reduction based on transcription factors confirmed the stable placode relationships relative to embeddings based on broader gene sets. When an UMAP was computed using the full expression profiles, placodal identities were partially overlapping in a specific order resembling anterior-posterior axial patterning. In line with this, the reclustering on transcription factors yielded a highly similar arrangement of placodal populations (Fig. 5g). This result supports the notion that transcription factor repertoires might be potentially enough to capture a compact, information-rich representation of placode identity and relatedness, and further motivates using DNA-binding protein sets for cross-taxon comparisons.

Finally, we used DNA-binding protein gene sets to infer similarity relationships among placodes and the *Ciona* preplacodal-like region using Cassiopeia. Although Cassiopeia was developed to reconstruct lineage trees from CRISPR “scar” character matrices, its parsimony framework applies to any discrete character table, including cross-species transcription factor repertoires. Here we discretized TF expression into binary characters (0/1 for absent/present), turning the comparison into a cladistic problem: identify the tree that explains the observed TF profiles with the fewest state changes. Cassiopeia searches for the topology that minimizes the total number of transitions across all TF characters and returns a parsimony cladogram in which placodes and species group by shared, evolutionarily parsimonious TF-expression patterns. In this setting, candidate cell-type homologies are evaluated by treating cell types (or structures) as “taxa” and conserved TF expression as “characters,” and asking whether the most parsimonious explanation consistently clusters corresponding types across species. In our case, both parsimony-based cladistic reconstruction with Cassiopeia and hierarchical clustering converged on a similar global organization (Fig. 5h): olfactory and lens formed the closest pair, adenohypophyseal-APF associated with this group, and trigeminal/otic/epibranchial formed a more distinct cluster. Notably, the *Ciona* preplacodal-like region was positioned adjacent to the olfactory-lens-adenohypophyseal-APF grouping (Fig. 5h), in line with the alternative method of gene usage overlap analysis and hierarchical clustering (Fig. 5f). Together, these results support a model in which vertebrate placodes draw on a deeply conserved DNA-binding protein repertoire, with especially strong similarity between olfactory, lens and adenohypophyseal programs, and with detectable continuity between vertebrate anterior placodal transcription factor usage and an anterior preplacodal-like regulatory state in *Ciona*.

Based on these results, we further examined the evolutionary origin of the olfactory placode from a complementary approach by leveraging single cell data and spatial transcriptomic maps (Fig. 6). Rather than restricting the comparison to placodes alone, we asked whether the olfactory placode shares a distinctive transcription factor repertoire with adjacent neural territories, as suggested by fate maps. This analysis revealed an unusually strong similarity between the olfactory placode and the most anterior forebrain domain that later produces the olfactory bulb, suggesting continuity within an integrated olfactory developmental program (Fig. 6a-c). Both regions expressed a shared set of anterior/neurogenic transcription factors, including *Vax1, Fezf1, Nr2e1, Otx1, Six6, Six3, FoxA2, Lhx2, Emx2*, and *Pax6*, among others (Fig. 6d and Fig. 7).

**Figure 6.**
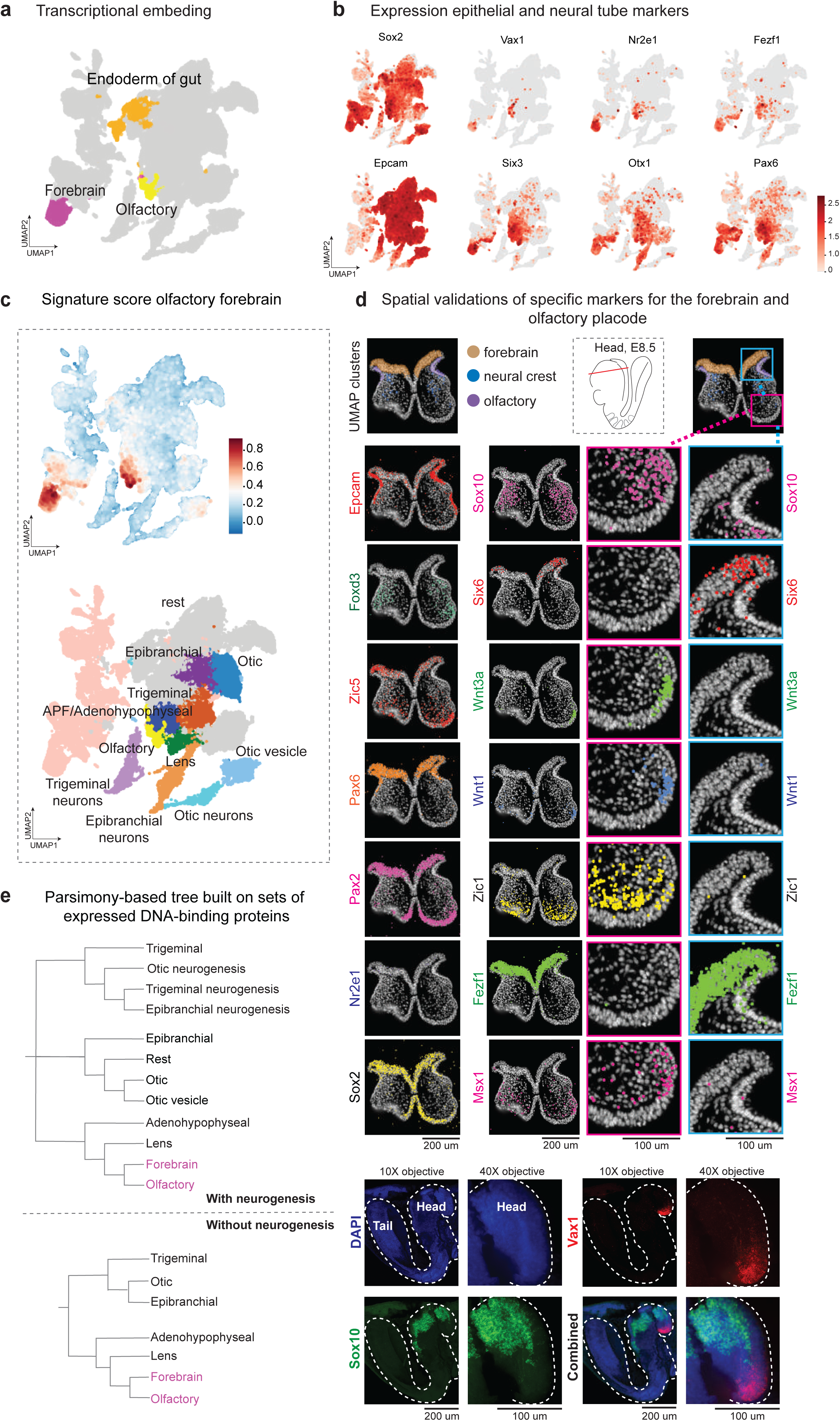
Similarity of transcriptional programs directing anterior forebrain and olfactory placode development. **(a)** Transcriptional embedding from Figure 1a-c, with highlighted forebrain (purple), gut endoderm (orange) and olfactory placode (yellow) highlighted. **(b)** UMAP feature plots showing expression of olfactory and forebrain markers. **(c)** Plotted gene expression signature showing commonly expressed genes specific for olfactory placode and anterior forebrain. **(d)** Spatial validations of specific markers for the anterior forebrain, neural crest and olfactory placode using the Xenium v1 platform at E8.5 of mouse embryo development. Bottom: HCR-based RNA *in situ* hybridization, DAPI nuclear labeling (blue), neural crest marker Sox10 (green), and olfactory placode marker Vax1 (red), 10X and 40X objectives. **(e)** Cassiopeia-built parsimony-based tree derived from a set of DNA-binding proteins expressed in cranial placode-containing clusters in relation to the anterior forebrain. The upper tree includes neurogenic derivatives of placodes. APF, anterior placodal field.

**Figure 7.**
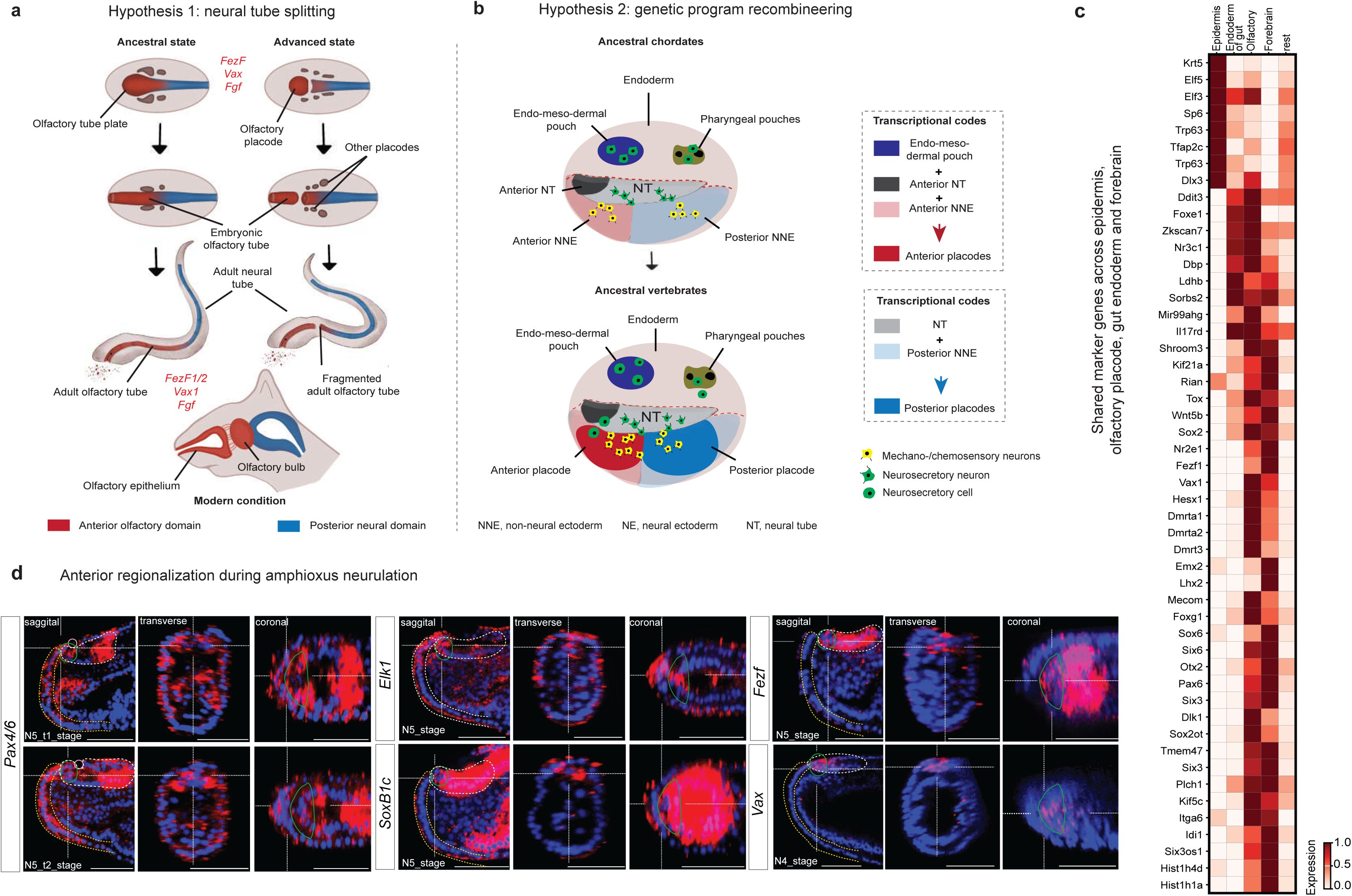
Single cell and spatial atlases help to refine the evolutionary origin of olfactory placode. (a-b) Evolutionary hypotheses proposing the origin of the olfactory placode. **(a)** The neural tube splitting hypothesis proposes that the tubular architecture of the nervous system originally evolved in basal deuterostomes in connection with olfactory function. According to this model, an ancestral tubular neuroepithelial structure underwent an anatomical partitioning that separated an anterior, olfaction-oriented domain from the remainder of the neural tube. The anterior portion subsequently gave rise to the olfactory placode, whereas the posterior segment developed into the centralized neural tube that later served as the core of the vertebrate nervous system^61^. **(b)** Hypothesis of olfactory placode evolution via transcriptional program recombination of endodermal, non-neural epithelial and anterior neural tube programs. In ancestral vertebrates, the recombined transcriptional codes gave rise to *de novo*-arising distinct anterior and posterior placodes, accompanied by an expansion of sensory and neurosecretory cell populations. **(c)** Heatmap showing shared gene expression in the anterior forebrain, olfactory placode, endoderm, and non-neural ectoderm. Populations ‘olfactory’, ‘forebrain’, ‘endoderm of nascent gut’ were identified using specific markers, while all other populations in the general UMAP embedding were grouped as ‘rest’. **(d)** HCR *in situ* hybridization assays showing the expression of anterior regionalization during amphioxus neurulation (N4-N5 stage). Green outlines the anterior neural plate border within the white dashed anterior neural tube; yellow dashes indicate the anterior surface ectoderm; pink outlines indicate the neuropore.

The spatial organization of the anterior neural tube area provides a developmental rationale for this similarity. At its rostral extent, the neural tube bordering the olfactory placode does not generate neural crest in consistency with the literature^45^, and placing the placode initially in direct apposition to prospective forebrain tissue. More laterally, however, neural crest migration brings crest-derived cells into proximity with the placode, creating a space where forebrain-adjacent and crest-associated signals can coexist. Importantly, this juxtaposition arises secondarily through neural crest migration rather than from an early adjacency of olfactory and crest-producing territories.

Together, these patterns suggest that the olfactory placode - more precisely, the anterior preplacodal domain that generates it - may be evolutionarily derived from an anterior neural-plate-like territory that became physically segregated from the neural tube. In this view, an ancestral anterior ectodermal/neural field could have been broader and more continuous, with the modern boundary between “neural plate” and “placode” emerging through tissue separation and physical reorganization rather than through an abrupt, functionally defined molecular program assembling *de novo* in evolution of chordate lineage.

To test whether this spatially observed affinity is recapitulated in an unbiased similarity reconstruction, we used Cassiopeia to build a parsimony tree from discretized DNA-binding protein expression profiles across placodal regions and the anterior neural tube. Consistent with the spatial transcriptomic comparison, this analysis placed the olfactory placode closest to the anterior forebrain (Fig. 6e), reinforcing the idea that olfactory placode development retains a particularly direct regulatory relationship to the anterior neural plate program, possibly being a result of a neural tube split into anterior and posterior portions in extinct chordates (the hypothesis is explained in Fig. 7a).

We also considered an alternative model proposed previously^46^, (explained in Fig.7b), in which the anterior placode is suggested to arise through a “blended” transcriptional program combining features of endoderm, non-neural ectoderm and anterior neural plate. To evaluate this possibility, we compared transcription factor repertoires between the olfactory placode, forming epidermis and endodermal territories in the spatial dataset. We detected only limited overlap in transcription factor expression, and the shared set was markedly smaller than the robust anterior/neural signature jointly expressed by the olfactory placode and anterior forebrain (Fig. 7c). Thus, within our data, regulatory similarity to endoderm and forming skin epidermis is weak relative to the strong affinity between the olfactory placode and anterior neural plate derivatives, suggesting that endodermal transcriptional programs blending could be limited during the elaboration of the olfactory placode program. However, that the quantitative degree of similarity is not necessarily important since not all gene recruitments will be equally important (a single, core TF may be able to import a whole battery of other genes, while others may be largely inconsequential).

Finally, we examined the expression of the transcription factors Vax, Pax6 and FezF in developing *Amphioxus* to test whether their domains extend from the anterior neural tube into adjacent ectoderm, as would be expected for a plesiomorphic chordate condition. In N5 larvae, all three genes are indeed expressed not only in the anterior neural tube but also in the anterior neural plate border region (Fig. 7d). Analysis of the *Amphioxus* single-cell atlas^47^, further confirmed their co-expression and broader transcriptional similarity between the anterior neural plate border and anterior neural tube, supporting the hypothesis that an ancestral anterior neuroectodermal field was later sub-divided into a future neural plate and an anterior ectodermal, sensory region.

Overall, our evolutionary analysis has a potential to reconcile two seemingly different origin models for the olfactory placode by pointing to a single underlying principle: an ancestral chordate embryo likely deployed a conserved anterior ectodermal neurogenic program that was not yet cleanly partitioned into “neural plate” versus “preplacodal ectoderm.” In vertebrates, neurulation and tissue reorganization would then have physically separated this once-continuous anterior field into two adjacent but developmentally distinct components of the olfactory system - the olfactory placode and the anterior forebrain that generates the olfactory bulb - while retaining substantial transcription factor continuity. This accounts for the unusually strong TF similarity we detect between olfactory placode and anterior forebrain in spatial transcriptomics, and it also explains why the *Ciona* anterior preplacodal-like ectodermclusters closest to olfactory (and lens) in cross-species TF-based comparisons: the *Ciona* domain can be interpreted as a persisting or redeployed version of the same ancestral anterior program, expressed in a different embryonic geometry. In this view, “*Ciona* preplacodal-like” and “anterior neural plate” are not competing ancestors but complementary readouts of a shared regulatory kernel, with the olfactory placode retaining a particularly direct - and thus potentially archetypal - connection to this ancestral anterior neurogenic module.

## Discussion

By combining single-cell transcriptomics, spatial transcriptomics and high-resolution clonal fate mapping, we provide an integrated view of how the vertebrate ectoderm is partitioned into neural tube, neural crest, placodal and epidermal territories. Our atlases help to refine the debated models of neural plate border specification^20,26^, which can differ between vertebrate taxa due to evolutionary adaptations to a general speed of embryo development, body size, presence of yolk, and heterochrony. In a strict “binary competence” model^30,48,49^, gastrula ectoderm is rapidly partitioned into a neural/neural crest-competent versus placode/skin-competent territory. By contrast, “gradient border” models posit that fate is specified stochastically according to local signaling environments^20,50^. In chick, many NPB cells occupy mixed states and resolve probabilistically into neural, neural crest, placodal, and epidermal fates, supporting the “gradient model However, in mouse, previous fate mapping data suggested that placodes and epidermis segregate from neural tube and neural crest from early stages^23^. Our new atlas reveals the basis for this organization: neural tube and neural crest form a closely connected module, whereas placodal cells lie within a broader continuum of Epcam⁺ ectodermal states. At E8.0-E8.5 we detect only a narrow “bridge” between the anterior placodal field and the neural/neural crest domain, which rapidly disappears, suggesting that the first major split at the neural plate border separates neural tube from a preplacodal/non-neural ectodermal sheet, with neural crest emerging later as a derivative of the neural tube rather than as a co-equal branch with placodes.

Within the ectoderm, we show that placodes do not arise as sharply segregated islands. Single-cell and spatial data instead support gradual border refinement from initially overlapping territories. At E8.5, placodes occupy distinct yet partially overlapping regions in transcriptional space aligned with their anterior-posterior positions. At each of these interfaces, we consistently identify “mixed” boundary cells co-expressing markers of neighboring placodes; similar mixed states occur where placodes meet prospective epidermis. Thus, the cranial ectoderm initially forms a continuous sheet with gradually emerging TF expression and fuzzy borders, from which placodal identities and boundaries emerge gradually via probabilistic or competitive resolution - providing a mechanistic basis for the clonal intermixing seen in our lineage data.

Our observations support a “competitive” model of placode assembly in which pre-placodal progenitors occupy broad epithelial territories and are progressively partitioned into adjacent placodes. Mixed-identity territories-cells co-expressing transcriptional signatures of neighboring placodes are particularly enriched at placode-placode and placode-non-placodal interfaces, and clonal data indicate that such cells preferentially give rise to immediately adjacent domains. Rather than presenting methodological or transcriptional noise, these intermediate states likely correspond to transient, bipotent or oligopotent progenitors whose fate is resolved by local signaling and mutual repression between competing GRNs. Spatial transcriptomics reinforces this view: boundaries between placodes are not molecularly sharp; instead, many placodal markers and positional genes display graded distributions, while a subset of border-enriched genes mark narrow transition zones. The observation that regulon activity domains are more compact than raw TF expression supports this notion of late, switch-like consolidation within broader, permissive fields.

Our NEPTUNE-guided TREX clonal atlas demonstrates that, despite extensive transcriptional overlap and mixed identities, clonal relationships tend to respect local spatial adjacency. However, a smaller fraction of cells contributes to combinations of neighboring domains - such as skin-placode, placode-crest, trigeminal-otic, and otic-epibranchial. The observations resulting from our clonal atlas are consistent with previous^34,35^ lineage tracing studies that highlight how clonal information reflects compartmentalization of the embryo^34,35^, and specifically agree with literature showing that the shared otic-epibranchial (posterior placodal) field contributes to both inner ear and epibranchial derivatives^51,52^.

Using SCENIC and STRING, we reconstructed GRNs for each mouse placode and found that each is driven by a coherent, largely self-contained combination of transcription factors and regulons that defines its identity and differentiation trajectory. At the same time, placodes share a substantial “combinatorial placodal toolkit”: overlap in differentially expressed genes is extensive, with especially strong overlap between olfactory and lens. Phylostratigraphy shows that DNA-binding proteins-based programs are dominated by evolutionarily old genes with a smaller lineage-specific contribution, supporting a model of combinatorial reuse and context-dependent reconfiguration of an ancient regulatory scaffold rather than placode-unique TFs. Regulon analysis along early neurogenic trajectories (trigeminal, otic, epibranchial) confirms early activation of classic ganglion TFs and reveals additional positional/signaling-related factors (e.g. Dlx3, Tbx3, Hoxc4), providing a roadmap for dissecting lineage-specific spatial modules. We note that these GRN models are based on co-expression, motif enrichment and literature-derived interactions; they necessarily contain false positives and omissions and should therefore be treated as a hypothesis-generating framework rather than a definitive regulatory map.

Our orthogonal ligand-receptor analysis shows that each placode expresses a characteristic set of numerous signaling ligands whose cognate receptors are found in neighboring placodes and adjacent neural tube, neural crest, and epidermis. Although correlative and awaiting functional validation, these patterns outline a rich network of potential short-range interactions, suggesting that placodal borders are shaped not only by intrinsic GRNs, global patterning gradients and signals from adjacent tissues such as cranial mesoderm, but also by local, placode-specific signaling that reinforces and sharpens emerging identities. Our cell-cell interaction predictions are based solely on co-expression of ligand–receptor pairs and should be considered as an outline of potential, rather than demonstrated, signaling relationships. Transcriptional presence of a ligand or receptor does not guarantee effective secretion, processing, binding or downstream pathway activation, which also depend on protein abundance, subcellular localization, post-translational modification and the presence of co-receptors and extracellular matrix components. The predicted networks therefore provide a useful catalogue of candidate interactions that may shape placodal borders and NPB patterning.

Taken together, our observations argue that, in the mouse, cranial placodes arise within a continuous, hierarchically patterned ectodermal field, where fuzzy borders, competitive resolution of mixed states, and locally tuned GRNs and signaling interactions jointly shape placodal territories. Within this unified framework, most placodes conform to shared organizational principles and exhibit extensive transcriptional and clonal interconnection, especially among posterior domains.

This view aligns with how placode homology is typically conceptualized. Placode homology is usually argued at the level of developmental origin (all cranial placodes arise from a shared preplacodal ectoderm adjacent to the neural plate), regulatory logic (deployment of a partially shared transcription factor toolkit), and morphogenetic/functional outputs (sensory/neurogenic derivatives that interface with the nervous system). In that framework, “homology” does not require that each placode is a direct one-to-one equivalent across distant taxa. Rather, placodes can be viewed as serial homologs within vertebrate-distinct modules produced by divergence from a common preplacodal program - and as deep homologs across chordates, where parts of the underlying gene-regulatory circuitry predate vertebrates and can be traced in more basal relatives.

Our results support this regulatory version of homology. In other words, divergence among placodes may occur largely downstream of, or in combination with, a shared regulatory scaffold rather than through completely unique transcription factors for each placode. Within that scaffold, the *Ciona* “preplacodal” population aligns most closely with the olfactory (and secondarily lens) placode at the level of shared DNA-binding proteins and inferred tree structure. A parsimonious interpretation is that the *Ciona* preplacodal region reflects an ancestral placode-like regulatory state - a proto-placodal/neurogenic module in the anterior ectoderm - that vertebrate later partitioned and specialized into multiple placodes. Under this model, the olfactory placode retains the greatest similarity to that ancestral state, either because it diverged less (conservation) or because it repeatedly reuses a core neurogenic patterning logic that is ancient within chordates. The consistent clustering of *Ciona* near the olfactory-lens group strengthens the argument that at least one vertebrate placode class - most plausibly the olfactory placode - sits closer to a conserved pre-vertebrate regulatory configuration than do other placodes. This leads to a testable “archetypical cranial placode” hypothesis: the olfactory placode may represent a basal placodal archetype from which other cranial placodes could be derived by adding lineage-specific regulatory layers (e.g., shifts in signaling responsiveness, cranial neural crest interactions, or placode-specific differentiation programs). Importantly, “most ancient” here should be framed carefully: it need not mean that olfactory placodes appeared first as an anatomical structure, but that their core regulatory circuitry may be closer to the ancestral chordate ectodermal neurogenic module detectable in *Ciona*.

When this comparative framework is extended across the entire cranial ectoderm, one structure emerges as distinctly divergent from the posterior placodal pattern: the olfactory placode. The olfactory placode exhibits developmental relationships and transcriptional affinities that depart from the patterns governing posterior placodes. Indeed, the olfactory placode is unusual among cranial placodes in several respects. First, its primary epithelial border is with the most anterior neural tube rather than with neural crest. Clonal reconstructions reveal essentially no shared progenitors between olfactory and posterior placodes (not surprising given the long distance between them), in stark contrast to the more prominent posterior placodal interconnection. Instead, olfactory clones are robustly shared with the adjacent non-neural surface ectoderm, and anterior neural tube derivatives. This indicates that olfactory progenitors straddle the boundary between surface ectoderm and neuroectoderm, but within a highly restricted anterior domain. Second, the most anterior neural tube region, which gives rise to the olfactory bulb and is contacting the olfactory placode, shares a significant portion of its transcription factor code (featuring *Vax1, Pax6, Otx2, Sox2, Six3, Nr2a1* and *Fezf1/2*, among other factors) with the neighboring olfactory placode. This shared GRN signature is not recapitulated in posterior neurogenic placodes or in more posterior forebrain domains, arguing against simple convergence. Functionally, both the anterior neural tube and olfactory system preserve epithelial organization and life-long epithelial neurogenesis. The olfactory epithelium contains radial glia-like progenitors, maintains neurogenic capacity throughout life, and harbors neurons that remain embedded within an epithelial context, paralleling features of neuroepithelium. Third, several specialized properties of the olfactory system blur the conventional boundary between CNS and PNS. For example, cells of olfactory epithelium can generate oligodendrocytes under experimental conditions, suggesting a latent CNS-like potential^53^. Gonadotropin-releasing hormone (GnRH) neurons are born within the olfactory epithelium and migrate into the forebrain, forming a developmental bridge across the ectoderm-neuroectoderm boundary^54–56^. Besides, other neurons residing in the chick brain were found to originate from the olfactory placode^57^. These observations are naturally accommodated if olfactory and anterior neural tube derivatives share a common ancestral epithelial neurogenic territory. Our Cassiopeia-based parsimony analysis of GRN similarity further supports this view. When placed in a broader phylogenetic context that includes tunicate single-cell data^54^, the mouse olfactory placode ends up being closest to the anterior proto-placodal field of *Ciona intestinalis* (in agreement with previous suggestions by Michael Levine and coauthors)^58^, and, within the mouse, closest to the anterior forebrain giving rise to olfactory bulbs. This arrangement is consistent with an evolutionary scenario in which an anterior proto-placode/neural plate territory, bearing an olfactory-like GRN, was present in early chordates and later resolved into separate neural tube and olfactory domains in vertebrates.

Comparative embryology lends additional weight to the unique developmental status of the olfactory placode. In cyclostomes, a single midline olfactory tube develops immediately anterior to the neural tube^59^, reminiscent of a continuous neural tube-like anterior neuroepithelial structure with olfactory specialization. Interestingly, a conceptually similar idea has recently been proposed for the evolution of the vertebrate visual system, suggesting that the ancestral bilateral eyes may have arisen through the duplication or separation of a single median eye associated with the neural tube midline^60^. Taken together, these findings suggest that the vertebrate olfactory placode did not emerge *de novo* from generic surface ectoderm but instead represents a domain that split off from an ancestral anterior neural-plate placode. In this framework, the early neural plate can be viewed as a large, continuous placodal field, with the olfactory system representing one of its most ancient sensory specializations. This perspective provides a developmental and evolutionary explanation for why the neural tube assumes a tubular architecture and how this geometry may relate to its ancestral chemosensory function. Building on this idea, our results support a previously suggested reinterpretation of the evolutionary origins of the epithelialized, lumen-bearing structure of the vertebrate CNS. This intriguing scenario is based on the assumption that early neural tubes enhanced directional olfaction by virtue of an enclosed, fluid-filled cavity that improved the capture, retention, and sampling of chemical cues^61^.

Another aspect of this model concerns the relationship between the olfactory-neural tube split and the origin of the neural crest. The *Vax1+* anterior neural tube domain adjoining the olfactory placode is, to our knowledge and in agreement with literature^62–64^, the only substantial region of the neural tube that does not give rise to neural crest cells. This spatial coincidence raises the possibility that the separation of an anterior olfactory territory from the rest of the neural plate predates, and perhaps constrained, the emergence of neural crest. Mechanistically, the absence of neural crest at the anterior neural folds and at the immediate neural tube-olfactory placode interface reflects a local suppression of Wnt signaling. When *Tcf7l1* is conditionally inactivated, this repression is lifted, allowing ectopic Wnt/β-catenin activity in the anterior neuroectoderm that reprograms these cells toward a neural crest fate^65^.

Overall, based on the above arguments, the olfactory placode retains a deep connection to the CNS-like ancestral state, both in its transcriptional program and in its clonal and morphological relationships. This helps explain why olfactory biology so often defies neat CNS/PNS categorization^57,66^, and suggests that certain “oddities” of the olfactory system - such as life-long epithelial neurogenesis, and migratory neuroendocrine neurons - are evolutionary remnants rather than uniquely evolved features.

Although our data support homology between the olfactory placode and the most anterior neural tube, alternative evolutionary scenarios must also be considered. Notably, one of us have proposed that anterior placodes may have arisen through partial co-option of endodermal and non-neural ectodermal, in addition to neural gene regulatory programs, including transcription factors such as FOXE1 and DLX3^46^. In our mouse datasets, however, we observe only limited overlap between endodermal, non-neural ectodermal and olfactory placode regulatory networks. While this does not exclude endodermal and non-neural ectoderm contributions, it provides only a limited support for this model than for the hypothesis of an ancestral olfactory-forebrain continuum.

At the same time, it is plausible that the olfactory placode ultimately became different from the olfactory brain via modifications of the GRN and recruitment of TFs, possibly from non-neural ectoderm and endoderm. Therefore, the two presented models are not incompatible. Indeed, some overlap between endodermal, non-neural and olfactory placode regulatory networks provides at least some support for such integrative view. Evolutionarily, recruitment of TFs from the non-neural ectoderm and anterior endoderm may have helped to ultimately separate the olfactory placodal territory from the anterior forebrain, thus allowing to reconcile the two evolutionary scenarios proposed.

Together, our data support a view of the cranial ectoderm as a hierarchically patterned field in which neural, neural crest, placodal and epidermal territories emerge by gradual resolution of fuzzy borders rather than precise early hard partitions. Within this framework, most placodes conform to shared organizational principles built on an ancient, reusable GRN scaffold, whereas the olfactory placode stands out as a special case that retains an unusually direct developmental, transcriptional and clonal connection to the anterior neural tube. This, in turn, is consistent with our hypothesis that an ancestral anterior neurogenic territory in early chordates was later split into forebrain and olfactory domains, with the olfactory system preserving key features of this archetypal epithelial neurogenic module and potentially constraining the emergence of neural crest. By providing a unified single-cell, spatial and clonal atlas, we offer both a developmental mechanism and an evolutionary framework for placodal diversification and olfactory origins, and we define a set of concrete GRN- and signaling-level predictions that can now be tested experimentally in vertebrate and invertebrate chordate models.

## Materials and methods

### Animal experiments

Animal research as part of this study was approved by the Ethical Committee on Animal Experiments (Stockholm North committee) and performed in compliance with the recommendations outlined in The Swedish Animal Agency’s Provisions and Guidelines for Animal Experimentation. (Permit number N15907-2019/18314-2021 and 19542-2024). Baylor College of Medicine Institutional Animal Care and Use Committee (Protocol#AN-4956). Laboratory animals were kept in standardized conditions (24 °C, 12:12 h light–dark cycle, 40–60% humidity, food, and water ad libitum). Animals from non-genetically modified mouse lines CD-1 and C57BL/6 were purchased from Charles River Laboratories and Janvier.

All Foxi3 mouse work was conducted according to NIH/ AVMA regulations and Institutional Animal Care and Use Committee policies at Baylor College of Medicine.

#### Generation of *Foxi3^CreER/+^;R26^tdT/tdT^* mice

Rosa26-tdTomato (*B6.Cg-Gt(ROSA)26Sor^tm^*^9*(CAG-tdTomato)Hze*^/*J*; stock 007909) mice were originally generated at the Allen Institute but procured from the Jackson Laboratory. To generate *Foxi3^CreER/+^;R26^tdT/tdT^* mice, *Foxi3^CreER/+^* studs were crossed to *R26^tdT/tdT^* females and selected and bred to carry both alleles. Foxi3-CreER mice (Foxi3^em2(cre/ERT2)Akg/J^ ^67^; are available from the Jackson Laboratory (strain #:038587). Heterozygous Foxi3-CreER mice were mated to homozygous ROSA-tdTomato reporter mice (B6.Cg-Gt(ROSA)^26Sortm9(CAG-tdTomato)Hze/J^ ^68^; available from the Jackson Laboratory. Wildtype (WT) embryos were used as negative controls in screening for tdTomato by fluorescence microscopy. Pregnant females were gavaged with two doses of 2 mg tamoxifen and progesterone dissolved in peanut oil at embryonic day 6.5 and day 7.5 (24 hrs apart).

### Single cell dissociation

C57BL/6J mice embryos were collected at embryonic day (E) 8.5 and E9.5. At E8.5, heads were isolated, whereas at E9.5 three regions anatomically were dissected: (1) olfactory; (2) lens, trigeminal, and adenohypophyseal; (3) otic and epibranchial regions. Samples were transferred to Eppendorf tubes containing 500 µL of 0.05% trypsin/0.02% EDTA (Sigma, 59417-C) and incubated at 37 °C for 15 min with gentle swirling and pipetting every 5 min. After 10–15 min, enzymatic dissociation was stopped by adding 500 µL of 10% fetal bovine serum (FBS) in PBS. Samples were centrifuged at 500 rcf for 5 min at 4 °C, followed by two washes with 500 µL of PBS containing 10% FBS. The supernatant was carefully removed, and the cell pellet was resuspended in 500 µL of HBSS supplemented with 1% FBS and kept on ice thereafter. The cell suspension was subsequently filtered into round-bottom polypropylene test tubes (Falcon) through a 40 µm cell strainer made of woven polyethylene terephthalate mesh (pluriSelect pluriStrainer, SKU 43-10040-40).

Foxi3^CreER/+^;R26^tdT/tdT^ embryos were harvested in sterile, cold 1X PBS and kept on ice until screened for Cre (tdTomato+). The head and neck of each embryo was dissected (up to the first somite at E8.5 and just above the heart at E9.5), and the samples were separated into wells by genotype. Embryonic tissue was next dissociated for 15-20 minutes in 0.22 µm papain/ DNase solution at 37 deg C. Following dissociation, tissue was homogenized by adding cold 1X CMF PBS + 2% FBS solution (1:1) and pipetting 100x per sample. Samples were centrifuged at 470 RCF for 5 min at 4 deg C, then supernatant was replaced with fresh 1X CMF PBS + 2% FBS solution. Sample pellet was re-suspended by pipetting. This process of centrifugation and resuspension was repeated a second time for more viscous samples. Finally, samples were filtered through 40 µm pluriSelect mini strainers into sterile 5 ml conical tubes.

### FACS sorting

Flow cytometry sorting was performed using FACSAria Fusion or FACSAria III instruments (BD) equipped with a 100 µm nozzle and a 1.5 neutral density filter, and controlled via FACSDiva software (BD). Wild-type mice lacking fluorescent reporters were used, and negative populations were isolated. Samples were maintained at 4 °C and gently agitated during sorting. The event rate was kept below 4,000 events per second. Debris, dead cells, and doublets were excluded based on forward- and side-scatter area versus width gating. Sorted cells were collected directly into low-binding tubes containing 50 µL of HBSS supplemented with 1% FBS.

Purified TDTOMATO+ cells from each sample were collected by FAC sorting in 150 µl fresh 1X CMF PBS + 2% FBS solution, using Cre negative (WT) embryos as a negative control for TDT+ cell selection.

### cDNA library preparation

The 10X Genomics protocol (Chromium™ Next GEM Single Cell 3’ GEM, Library & Gel Bead Kit v3.1) was used to prepare cDNA libraries for each set of single cell RNA-seq experiments using C57BL/6J and Foxi3^CreER/+^;R26^tdT/tdT^ mice embryos. In summary, Gel Beads-in-emulsion (GEMS) were produced via mixing Single Cell 3’ v3.1 Gel Beads with the cell solution master mix for each sample and Partitioning Oil, then loading these into the Chromium Chip G. Lysed gel beads were combined with Illumina TruSeq R1 primers, a 12 nt unique molecular identifier (UMI), 16 nt 10X Barcode, and 30 nt poly(dT) sequence for each GEM, as well as a reverse transcription Master Mix of reagents. After generating pooled, barcoded full-length cDNA from the poly-adenylated mRNA of each cell, silane magnetic beads were used to purify cDNA from this mixture, then PCR was used to amplify the cDNA. Appropriately sized fragments were selected for the library, and paired end P5 and P7 primers, the TruSeq R2 primer sequence, and a sample index were added subsequently.

### Single cell RNA-sequencing & mapping to Mus musculus reference genome

The de-multiplexing of scRNA-seq data for all experiments was performed by Novogene, subsequent to sequencing the cDNA. 10X Genomics software Cell Ranger v8.0 (https://www.10xgenomics.com/support/software/cell-ranger/8.0) was used for barcode-processing, gene counting/ aggregation, and sequence alignment to a reference genome. The GRCm38 Mus Musculus NCBI reference genome was modified to include the tdTomato-WPRE sequence for mapping the tdTomato gene in data processed for analysis.

### Single-cell RNA-seq datasets and quality control

Single-cell RNA-seq datasets from Foxi3-traced and wild-type embryos at embryonic days 8.5 (E8.5) and 9.5 (E9.5) were processed in Scanpy (v1.9.8)^69^. For each of the 15 datasets we excluded low-quality cells based on gene detection, UMI counts, and mitochondrial transcript fraction. Doublets were predicted using Scrublet (v0.2.3)^70^ and removed — with doublet rates ranging from 0.1 % to 7 % per dataset. Quality-control metrics for all samples are summarized in Quality control (Supplementary Table 4).

### Data integration and normalization

After filtering, all datasets were merged, using concatenate (join=’outer’). Resulting in an object with 62,269 cells and 25,865 genes. Expression values were normalized to 10,000 counts per cell and log-transformed^71^. 2000 highly variable genes were identified for downstream analyses using Scanpy v1.9.8.

### Dimensionality reduction, clustering and annotation

To capture cell expression similarity, PCA with 30 components was used to reduce dimensionality and construct a k-nearest-neighbor graph (k=30 neighbors). Cell populations were identified using Leiden clustering^72^, (resolution = 1.2) and visualized with UMAP^73^. The top 50 cluster markers were detected using the Wilcoxon rank-sum test^74^, and major populations—epithelial (EPCAM⁺), mesenchymal progenitors, neural crest, neural tube, notochord, and vascular lineages—were annotated using established marker profiles.

### EPCAM^+^ sub-setting and selecting placodal populations

To focus on placodal populations, we computationally isolated EPCAM-positive epithelial clusters (≈43,000 cells) and analyzed them independently. After re-normalization, log-transformation and highly variable gene selection (using default parameters), PCA was recomputed with 40 components. Clustering employed a neighbor graph with k=30 neighbors, followed by Leiden clustering (resolution = 0.6) and UMAP embedding. Clusters in the EPCAM subset were annotated and marker genes visualized using the same strategy as used for the full dataset, identifying sixteen epithelial cell types, including six distinct placodal populations.

### RNA velocity and developmental trajectory analysis

RNA velocity analysis on the EPCAM-positive cells was estimated using scVelo (v0.2.5)^75^. Spliced and unspliced transcript counts were first quantified with Velocyto^76^ from CellRanger-aligned BAM files. Local transcriptional transitions were modeled using 3,000 highly variable genes, with moments computed across 30 neighbors and 40 components in PCA space. Velocities were estimated using a deterministic model, and the velocity vector field was visualized by projecting streamlines onto the existing EPCAM UMAP embedding.

### Placodal lineage clustered based on a set of DNA-binding proteins

To explore cranial placode lineages, we computationally isolated placodal clusters (otic, trigeminal, epibranchial, olfactory, lens, and adenohypophyseal) and reclustered cells using DNA-binding transcription factors as features. After normalization, PCA (35 components) was used to reduce dimensions, and cell groups were defined using Leiden clustering on a 25-neighbor graph in PCA space. Lineages were visualized with UMAP, and clusters annotated using well-established transcription factor expression profiles.

### Comparison with Ciona intestinalis

*Ciona intestinalis* forms a common preplacodal epithelial region instead of separate cranial placodes, suggesting this tissue may reflect the ancestral origin of chordate placodal programs. To compare this domain with our mouse placode data, we analyzed publicly available single-cell RNA-seq cionas datafrom GEO (accession GSE131155). Raw gene-by-barcode matrices for each sample were downloaded in HDF5 format and processed using Scanpy (v1.8.2) in Python 3.8. Gene expression matrices were merged using concatenate (join=’outer’). Following matrix integration, cells expressing less than 300 genes or more than 6,500 genes were removed, as these profiles most likely represented low-quality cells or doublets. The remaining dataset was normalized to10,000 counts per cell, log-transformed, and projected to reduced dimensions by PCA (35 components). Cell groups were identified by Leiden clustering on a 45-neighbor graph in PCA space (resolution = 1.3) and visualized using UMAP. The preplacodal region in the Ciona intestinalis dataset was identified by expression of marker genes (SIX1, SP8, FOXG1/FOXI1, ISL1). These marker profiles were visualized in UMAP space.

### Phylostratigraphy analysis

For the phylostratigraphy analysis, we used the phylogenetic framework developed by Šestak and colleagues^41^, which comprises 20 phylostrata. Each phylostratum corresponds to a major evolutionary transition, spanning from the Last Common Ancestor (LCA) of all cellular organisms to Mus musculus.

### Parsimony-based TF program reconstruction

To identify transcription factors shared between Ciona preplacodal and mouse placodal groups, each cluster was analyzed independently. Genes detected in ≥7% of cells were selected in both the epithelial (EPCAM-positive) and Ciona datasets, and intersected with a published reference list of mouse transcription-factor DNA-binding proteins^77^. TF detection across placodes was summarized in a binary TF-by-placode dataframe (1 = present, 0 = absent). TFs detected in all or none of the placodes were removed to keep only informative, lineage-defining shared TFs. Shared TFs were then identified using pairwise overlap analysis between each mouse placode and the Ciona preplacodal cluster. Lineage relationships were reconstructed using Cassiopeia (v2.0)^78^. Parsimony trees were solved using integer-linear programming (convergence limit = 500, graph layer cap = 500, unweighted, seed = 1234) on the binary TF dataframe described above.

### Placodal lineage analysis (epibranchial, trigeminal, and otic)

Each placodal lineage was analyzed independently using DNA-binding proteins as the feature set. All subsets were scaled, reduced by PCA (35 components). Cell neighborhoods were defined using a kNN graph (25 neighbors in PCA space), and clusters were assigned with Leiden. Lineage and cluster relationships were visualized in UMAP space.

### Trajectory reconstruction with scFates

Trajectories were reconstructed using scFates v0.4.1 (Faure et al., 2023) by fitting an ElPiGraph in UMAP space (55 nodes, μ = 50, λ = 0.01), generating a soft cell–node matrix (0–1). We set the root manually using early placodal clusters and calculated pseudotime as the geodesic distance along the graph. Pseudotime-associated TFs were detected (association threshold A ≥ 0.5), grouped into transcriptional modules using a 60-neighbor PCA graph and Leiden resolution 0.4, and visualized as scFates pseudotime heatmaps.

### SCENIC regulon inference

Gene-regulatory networks were built with pySCENIC (v.0.12.1)^79^, for each placodal subset using raw count matrices. TF–target co-expression links were scored with GRNBoost2 and refined using cisTarget with default parameters. Regulon activity was quantified per cell, producing a TF-by-cell activity matrix. We then projected regulon signals onto the existing UMAP embeddings to visualize regulatory patterns within each lineage.

### Cell-cell interaction analysis

Cell-cell communication among placodal, neural crest, neural plate, and epithelial populations was calculated in Python using LIANA v0.1.9^80^. Ligand–receptor interactions were inferred with CellPhoneDB v.2.0^81^ (expression threshold = 0.35).

### Placode gene regulatory networks

We used pySCENIC (v0.12.1)^82^ to infer separate gene regulatory networks for the early (E8.5) and late (E9.5) placodal cells in our transcriptomic atlas. The early GRN encompassed highly expressed genes and cells from the adenohypophyseal, epibranchial, otic, and trigeminal placodes, and the late GRN encompassed cells and genes from the olfactory and lens placodes. For both GRNs, we used ‘rank_threshold=1000’ and ‘auc_threshold=0.01’ to identify high-confidence targets of enriched motifs from the cisTarget mouse database (Imrichová, 2015, NAR). We used ‘top_n_regulators=20’ to generate the final TF-target gene modules and identify the top regulatory hubs in each GRN.

### STRING interacting protein network analysis

Protein–protein interaction networks were generated using STRING v12.0 (https://string-db.org/) with a minimum confidence score of 0.7 for olfactory placodes and 0.5 for the remaining cranial placodes. The list of transcription factors (TFs) used in the analysis is provided in Supplementary Table 3.

### Tissue preparation for Xenium spatial transcriptomics

C57BL/6J mouse embryos were collected at E8.5 and E9.5 and fixed overnight in 4% PFA. After fixation, the embryos were washed three times in DPBS, then dehydrated sequentially in increasing concentrations of ethanol (70%, 80%, 95%, and 100%), with 1 h incubation at each step. Following dehydration, samples were cleared in xylene and embedded in molten paraffin. One block contained approximately 13 E8.5 embryos, and the second block contained 11 E9.5 embryos. These blocks were used to generate sections according to the Xenium protocol. Tissue sections were cut at a thickness of 5 µm and placed onto a Xenium slide, followed by fixation and permeabilization. A total of 347 genes were included in the analysis. Gene panel probe hybridization was performed overnight (18 hours) at 50°C. Slides were washed the following day to remove unbound probes. Ligase was then added to circularize the paired ends of bound probes (2 hours at 37°C), followed by enzymatic rolling circle amplification (2 hours at 30°C). The slides were washed in TE buffer before background fluorescence was chemically quenched. Image processing, decoding, quality score generation, and DAPI-based cell segmentation were automatically performed using the Xenium Analyzer system. Data were acquired with instrument software version 2.0.1.0 and analyzed using Xenium software version 2.0.0.10.

### Data preprocessing, clustering, embedding and cell annotation of Xenium data

Each Xenium slide was processed independently in Python v.3.10 using Scanpy v1.10 and Squidpy v.1.4^83^. Raw outputs (feature matrix, cell metadata, spatial coordinates) were loaded into AnnData objects. To keep only high-quality cells, data from all slides were filtered using the following thresholds: 20–100 detected genes and 20–470 total transcripts per cell. Filtered data were normalized, log-transformed, and reduced in dimensionality using PCA with default parameters applied per slide. Slide-specific neighbor graphs were generated separately, using spatial parameters defined per slide: slide 1 (n_neighbors = 35, n_pcs = 15), slide 2 (n_neighbors = 30, n_pcs = 30), and slide 3 (n_neighbors = 45, n_pcs = 25). For each slide, we generated low-dimensional graph embeddings using UMAP and detected clusters with the Leiden algorithm (resolution = 6.5). Slide annotations were assigned independently by integrating spatial cell locations with cluster-level gene expression signatures.

### Reclustering of Epcam^+^ cell populations in Xenium data

Similar to single-cell analysis, we extracted only EPCAM-positive clusters from each slide and re-analyzed them as independent subsets using a uniform workflow: normalization, log-transformation, dimensionality reduction with PCA (90 components), neighbor graph construction (n_neighbors = 45, n_pcs = 80), and clustering using the Leiden algorithm (resolution = 2.5). Cluster-level annotations were assigned per subset, following the same approach used for the full dataset.

### Placodal and neural crest subset identification in Xenium data

To investigate trajectories shared among placodal populations, placode-associated clusters were selected from the EPCAM-positive embedding for each slide and analyzed as independent subsets. Specifically, for each spatially selected placodal region, neighborhood graphs were constructed using the same parameters across all subsets (*n_neighbors* = 15 on the X_spatial representation). Diffusion maps and diffusion pseudotime were computed independently for each subset, with root cells manually specified based on spatial position and marker expression. Cells were ordered along pseudotime, and pseudotime-aligned expression matrices were generated for curated panels of placodal marker genes. Heatmaps were then plotted for each subset to visualize gene expression dynamics along pseudotime. To analyze trajectories shared between placodal and neural crest cells populations, placodal and neural crest cells were extracted and analyzed together using the same workflow and parameter settings described above.

### Linearization of spatial coordinates along epithelial structures in spatial data

As many of our regions of interest followed an epithelial structure, we developed a tool to linearize the spatial coordinates of cells and transcripts along the epithelium and align the resulting spatial coordinate it to boundaries between cell types. First, regions of interest were selected in the Xenium Explorer 4 software and cell coordinates were exported. Cell coordinates and transcript coordinates were then loaded in R (v 4.3.2) and filtered for which belonged to the region of interest. Then, the entire set of cell coordinates were supplied together with a starting point and an end point, and a custom code script in R traced the outline of the region and aligned each coordinate to the progress along this outline. If a specific point of interest was present in the region (such as a border between two clusters of cell types), this point could be demarcated as position 0, and the coordinate was measured as a distance from this point. The output coordinate could then be used for plotting purposes as a linearized spatial variable. Figures were prepared using ggplot2. The necessary R libraries were vroom^84^ for importing the large transcript tables, dplyr^85^ and reshape^86^ for data handling, and ggplot2^87^ for plotting.

### Whole mount in situ hybridization in mouse embryos

The procedure was conducted in accordance with the Molecular Instruments v3 version of the protocol. (https://files.molecularinstruments.com/MI-Protocol-RNAFISH-Mouse-Rev11.pdf). C57BL/6J mouse embryos were fixed in 4% paraformaldehyde (PFA; Histolab) at 4 °C for 3 h. Following fixation, samples were washed in phosphate-buffered saline containing 0.1% Tween-20 (PBST) and dehydrated through graded methanol/PBST solutions before storage at −20 °C. Prior to analysis, embryos were rehydrated and permeabilized with proteinase K (10 µg ml⁻¹) for 15 min at room temperature, followed by postfixation in 4% PFA and additional washes in PBST. For Multiplexed HCR™ RNA-FISH, embryos were equilibrated in probe hybridization buffer, pre-hybridized, and incubated with probe mixtures overnight at 37 °C. Excess probes were removed using probe wash buffer and 5× saline–sodium citrate buffer supplemented with 0.1% Tween-20 (5× SSCT). For signal amplification, embryos were pre-incubated in amplification buffer and then incubated with hairpin amplification solutions overnight at room temperature. After washing with 5× SSCT, samples were imaged using Nikon Ti2E microscope with a Yokogawa Spinning Disk Confocal Unit.

### Ultrasound-guided microinjections of the barcoded lentivirus

Lentiviral injections were carried out following established methods^36^. Ultra-sharp glass capillaries were prepared by pulling and grinding. At E7.5, pregnant females were anesthetized with isoflurane, maintained on a heated surface, and given preoperative analgesia (buprenorphine, 0.05–0.1 mg/kg, subcutaneous). After sterilization and abdominal preparation, the uterus was gently exteriorized into a Petri dish containing PBS. Embryos were visualized using a Vevo2100 ultrasound system, and 46 nL of lentivirus was injected through the uterine wall into the amniotic cavity. The uterus was then returned to the abdomen, muscle incisions were closed with absorbable sutures, and the skin was sealed with sutures or staples. Animals recovered on a heating pad under supervision and received postoperative analgesia. Embryos were harvested at E11.5 following maternal euthanasia by CO₂ overdose and cervical dislocation and were subsequently euthanized by decapitation.

### Single cell dissociation for clonal experiments

Embryos were harvested at E11.5 following maternal euthanasia by CO₂ overdose and cervical dislocation and were subsequently euthanized by decapitation. Mouse embryos were dissected from the uterus, sacrificed, and transferred to ice-cold Hank’s Balanced Salt Solution (HBSS). Fluorescent reporter expression (tdTomato) was assessed using a fluorescence stereomicroscope. Embryos were dissected into different regions containing olfactory and forebrain; eye, trigeminal ganglia and midbrain and hindbrain; otic and epibranchial ganglia region; and the entire cranial region. Dissected tissues were placed in dissociation buffer containing Collagenase P (2 mg/mL) (Roche, Sigma-Aldrich), TrypLE (1×), and DNase I (1 µL/mL) prepared in HBSS with calcium and magnesium. Samples were then incubated for 20 minutes at 37 C in a preheated shaking incubator set at 150 RPM. After tissue were dissociated to stop the enzymatic were aded ice-cold HBSS supplemented with 2% FBS. Cells were pelleted by centrifugation at 500 × g for 15 minutes at 4 °C, resuspended in HBSS containing 2% FBS, and kept on ice. Prior to downstream applications, single-cell suspensions were filtered through a 40 µm cell strainer. Flow cytometry sorting was performed using FACSAria Fusion or FACSAria III instruments (BD) equipped with a 100 µm nozzle and a 1.5 neutral density filter, and controlled via FACSDiva software (BD). Single-cell suspensions were processed for library preparation according to the 10X Genomics protocols described above.

### scRNA-seq data preprocessing for clonal experiments

Settings and code used for processing are available on this project’s GitHub repository. See Code Availability Statement. After 10x Genomics Chromium sequencing of samples, the reads were aligned using 10x Genomics Cell Ranger (v.7.0.1) [Source: Cell Ranger] with chimeric mm39 and FP genome reference. The TREX v.0.4.dev402^38^ workflow was used for extracting clonal barcodes and reconstruction of cells which generate cloneID count matrices from the output of Cell Ranger. Initial quality control and preparation of data was used using scanpy (v.1.10.3)^69^ and custom python (v.3.11.10) functions. Cells were filtered for gene expression count, percentage mitochondrial UMIs, and number of genes by counts. Barcode correction was performed for each relevant cloneID. Clone labels were set for cells for each sample, with a minimum clone size of 2, resulting in a cloneID annotated gene expression object. Doublet detection and filtering were performed using scanpys built in Scrublet method^70^, using thresholds for each sample. Cells with multiple clone ids were considered doublets and subsequently filtered out. Dissected samples for placode areas and full cranial dissections were integrated separately and then together into one object. This process included concatenation of objects, followed by filtering genes with counts below 5. Counts were then normalized to have a total count of 10 000 per cell and then log-transformed (Ahlmann-Eltze andHuber 2023). Highly variable genes were identified, and the top 5000 were selected for scaling followed by PCA. The top 30 PCs were used for integration using Harmonypy (v.0.0.10)^88^ in symphonypy (v.0.2.1)^89^. Using the PCs of the integrated object, a nearest neighbor distance matrix was computed using a k=30 which was used to create a gene expression UMAP embedding of the integrated object with umap-learn (v 0.5.7)^90^. Cells were clustered using the Leiden algorithm with leidenalg (v.0.10.2)^72^.

A similar process was used for creating subsets of the data. Desired cells were selected, and all other cells were masked from the concatenated object. Cell cycle genes^91^ and genes present in less than 3 cells were removed. See supplementary data for cell cycle genes. Cells with less than 100 expressed genes and Scrublet predicted doublets were removed. Afterwards the object was integrated following the procedure above starting from normalization.

### Cell type annotation in scRNA-seq clonal data

Cells were annotated by first calculating the differentially expressed genes of Leiden clusters using Welch’s t-test in CELLxGENE python implementation version cellxgene-(1.1.2). Then annotations for cells were defined based on if differentially expressed genes were visually identified in tissue of E11.5 mice using the Allen Brain Atlas or by description in literature. This process was repeated for different subsets of the data. Full annotations can be in supplementary data.

### Clone2vec machine learning framework for clonal embeddings and their analysis

Clonal clusters were computed using the clone2vec algorithm using sclitr (v. 0.1.4)^35^. To run clone2vec a nearest neighbor algorithm computed the top 5 nearest clonally labeled cells for each clonally labeled cell on the integrated dataset. Clone2vec was run using 50 epochs which outputs the resulting clone embedding. PCA was performed on the clonal embedding, and the top 20 PCs were selected for neighborhood detection and visualization. To construct a kNN-graph and UMAP based on latent representation of the clones’, silhouette scores^92^ for different k was calculated and k=10 was chosen as the appropriate number of neighbors. Clones were now visualized in the UMAP embedding and clone frequencies in clonal clusters calculated.

### Amphioxus husbandry and in situ hybridization of amphioxus embryos

Amphioxus husbandry, spawning induction were performed as described previously^47^. Whole-mount in situ hybridization of amphioxus embryos was performed as described in Bozzo et al., 2020^93^. Embryos were imaged using a Leica SP8 confocal microscope with Leica Application Suite X (LAS X) v3.5.7.23225, and images were processed using Fiji (ImageJ).

## Data availability section

The original datasets are deposited to GEO with the accession number, GSE317146 (single cell transcriptomics), GSE317114 (spatial transcriptomics), GSE318565 (clonal single cell data).

The computational notebooks are deposited to GitHub. Single cell and spatial transcriptomics analysis https://github.com/fateevajulia/Neurogenic-placodes-illumination-specification-and-evolution

Analysis of gene-regulatory networks for placode development https://github.com/calebclayreagor/placode-grn

Clonal analysis https://github.com/FelixWaern/clonal-placodes-2026

## Acknowledgements

I.A. was supported by the ERC Synergy grant (KILL-OR-DIFFERENTIATE), Swedish Research Council, Paradifference Foundation, Cancer Foundation in Sweden, Knut and Alice Wallenberg Foundation, ALSF (the Crazy 8 project grant), Austrian Science Fund (consortium SFB F78 program and Emerging Fields “Brain Resilience” consortium, as well as stand-alone project grants). A.K.G. was supported by NIH R01 DC013072. H.R.M. was supported by NIH F31 DE032898. A.S. was funded by the BBSRC: BB/V006290/1 and BB/V006339/1 grants. We acknowledge the In Situ Sequencing Facility at SciLifeLab, funded by Science for Life Laboratory and the Swedish Research Council, for providing in situ sequencing services, National Genomics Infrastructure, Biomedicum Flow Core, Biomedicum Imaging Core, Biomedicum Comparative Medicine for access to their facilities and services during the project.

## Author contributions

I.A., A.K.G., G.S., A.M., Y.F., F.W., H.R.M., A.T., C.C.R., and J.B. contributed to conceptualization and manuscript writing. I.A., A.K.G., A.M., Y.F., F.W., K.F., A.G.E., A.S., L.M.K., F.S., L.M.K., F.S., A.K., P.K., E.R.A., B.S. and S.I. performed the investigation. Y.F., F.W., H.R.M., A.T., S.I., C.C.R., J.B., K.A., and A.M. carried out computational analysis. A.M., Y.F., F.W., H.R.M., A.T., J.B., Z.K., K.F., I.K., B.S., A.G.E., A.K., L.M.K., F.S., and P.K. contributed to validation. J.B., I.K., and Z.K. generated high-content microscopy data. I.A., A.K.G., G.S., A.S., E.R.A., A.M., Y.F., F.W., K.A. and A.G.E. performed data annotation. I.A., A.K.G., and A.S. acquired funding and resources. I.A., A.K.G., A.M., Y.F., and F.W. supervised the study and managed project administration.

