## Supplementary Figures 1-17 for "Single-cell, clonal and spatial atlases of cranial placodes illuminate their specification and evolution"

### Supplementary Figure 1

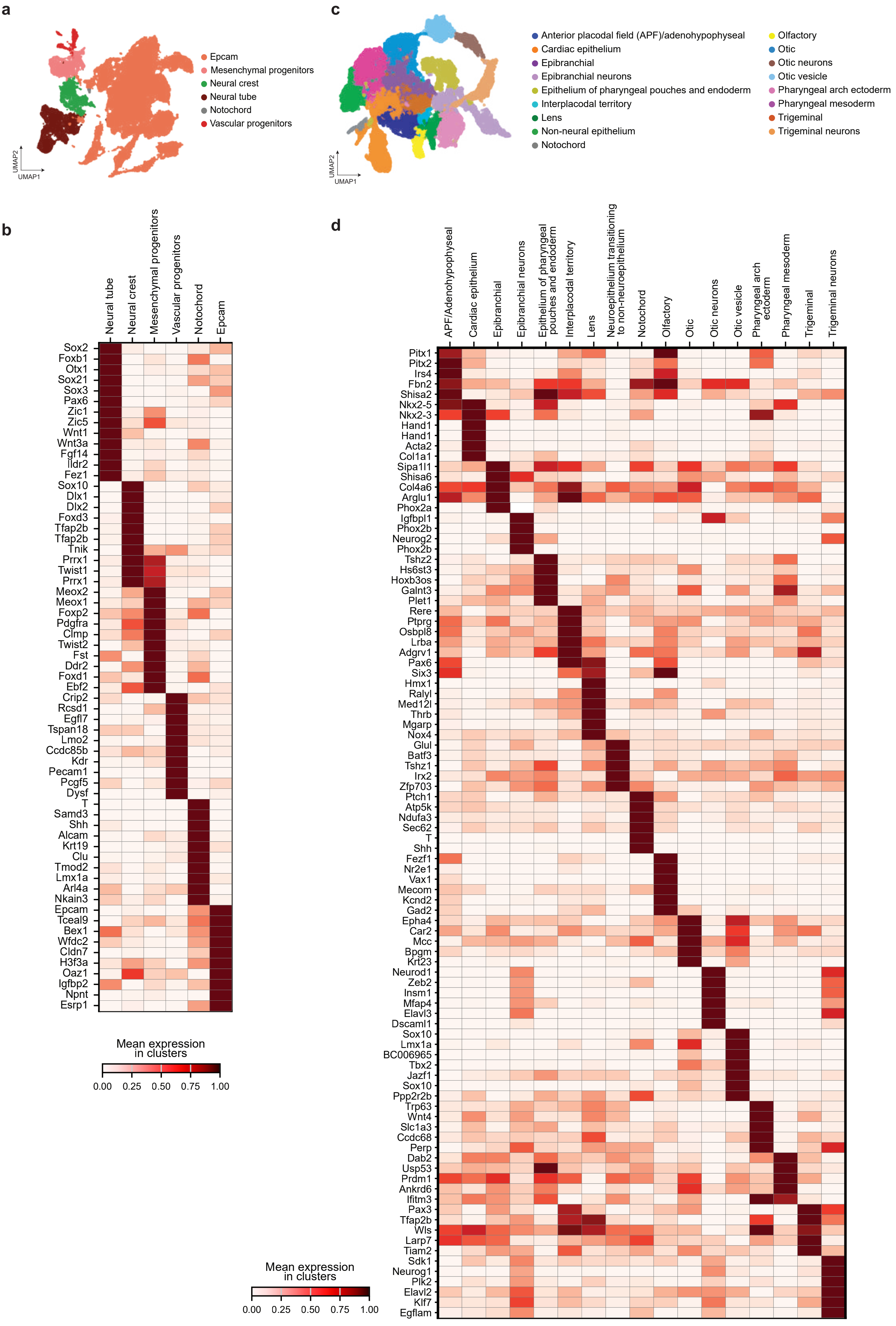

**Supplementary Figure 1. Transcriptomic profiling of Foxi3-traced and wild-type embryos at day 8.5 and 9.5 reveals distinct gene expression patterns and cellular heterogeneity.**

**(a)** UMAP embedding showing all clusters of ectodermal and mesodermal derivatives. **(b)** Heatmap showing the differentially expressed genes in ectodermal and mesodermal derivatives. **(c)** UMAP embedding showing annotation for the corresponding clusters. **(d)** Heatmap showing the differentially expressed genes in Epcam+ and placodal epithelia-derived cell populations.

Supplementary Figure 2

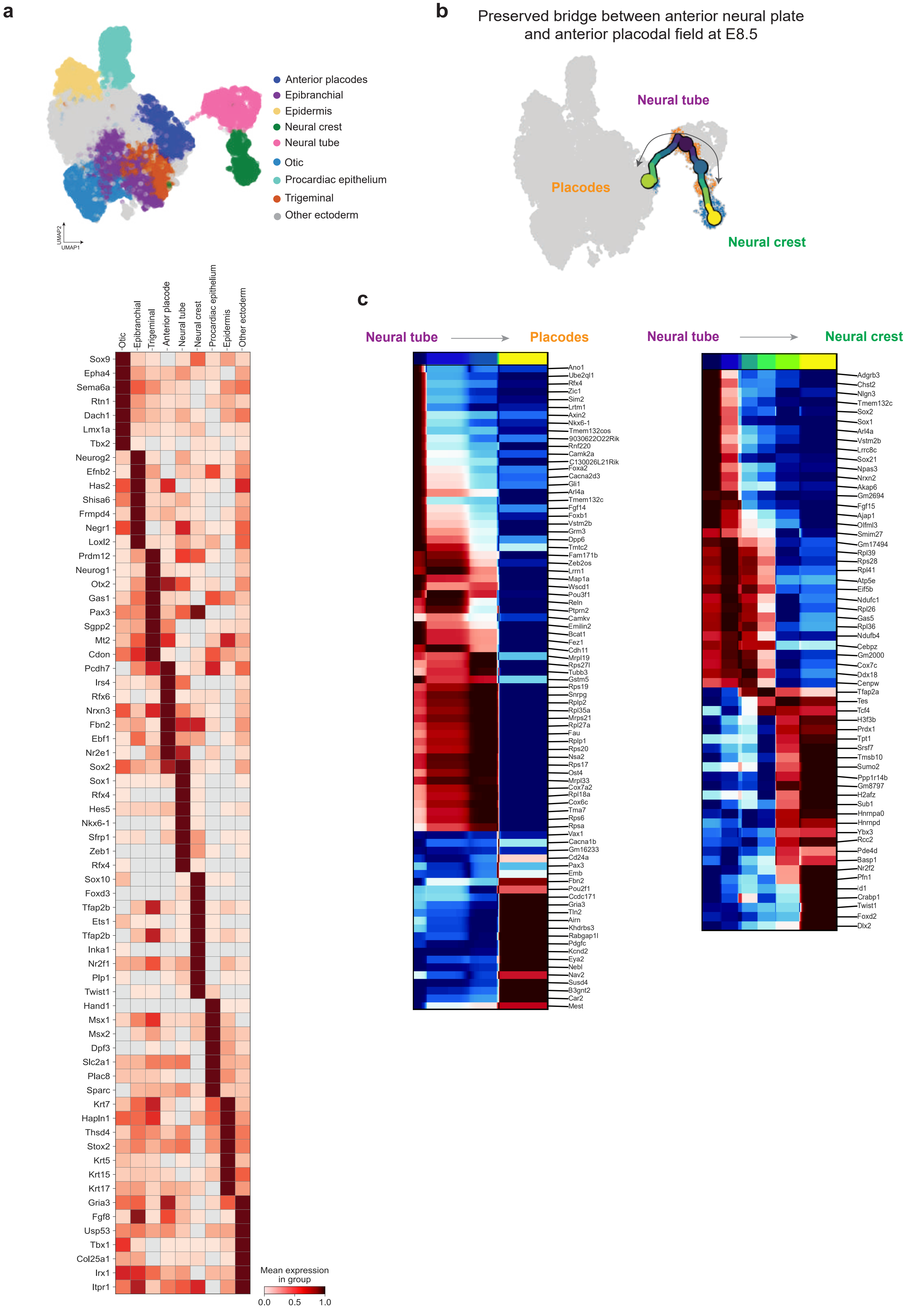

**Supplementary Figure 2. Clustering and trajectories of neural plate border derivatives form mouse embryos at day 8.5. (a)** Top, UMAP visualization and annotation of merged Foxi3 traced and wild-type datasets at embryonic day 8.5. Bottom, heatmap showing mean expression of selected genes across separate placodes, epidermis, neural crest, neural tube, procardiac epithelium and other ectoderm. Color scale indicates mean expression level within each cluster. **(b)** Preserved bridge between anterior neural plate and anterior placodal field at E8.5 **(c)** Pseudotime-integrated heatmaps of placode-to-neural tube-to-neural crest connection.

Supplementary Figure 3

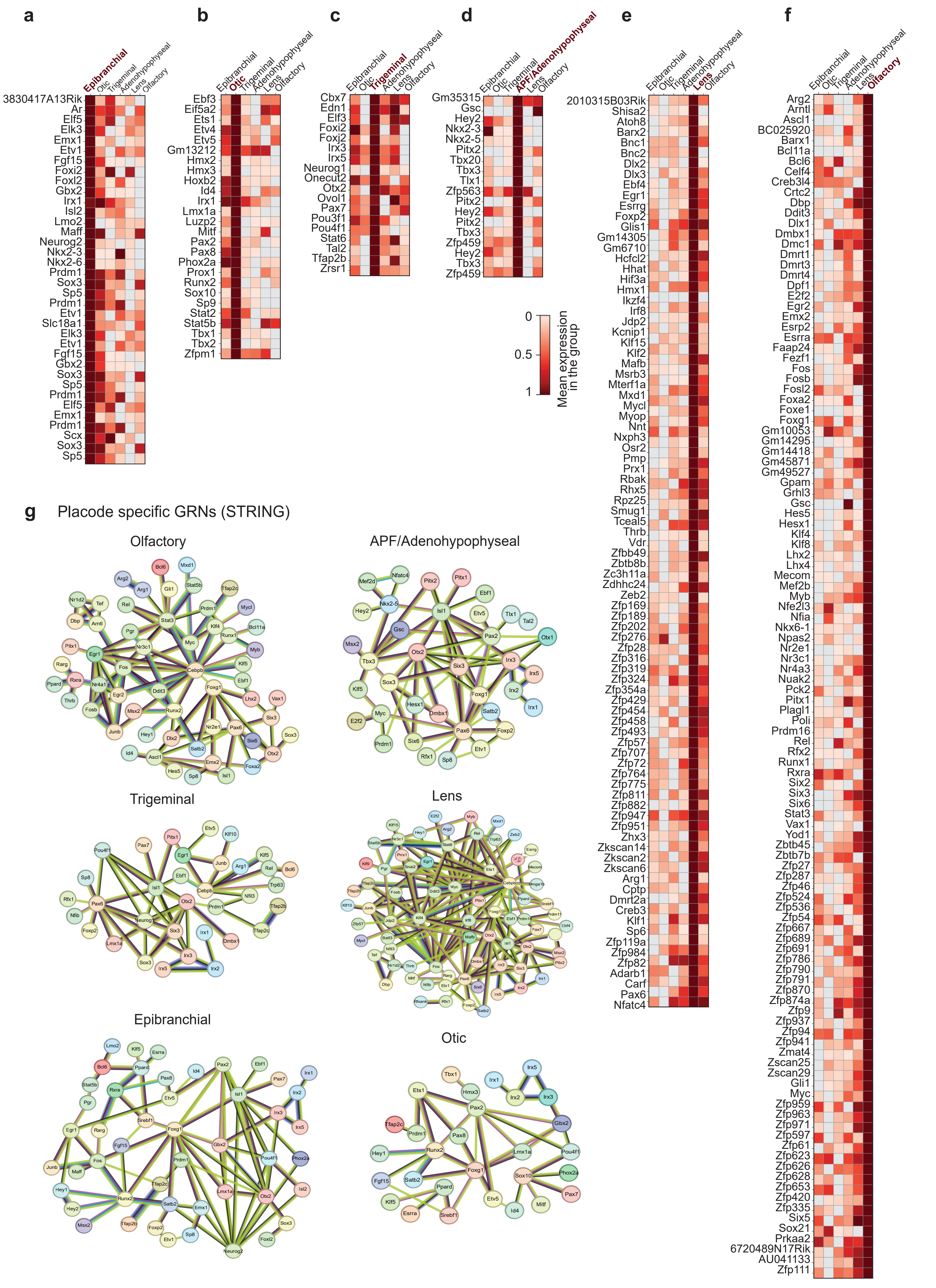

**Supplementary Figure 3. Differentially expressed DNA-binding proteins in clusters containing cranial placodes.**

**(a)** Epibranchial. **(b)** Otic. **(c)** Trigeminal. **(d)** Adenohypophyseal. **(e)** Lens. **(f)** Olfactory. **(g)** STRING-based connections of GRN components highlighting possible interactions of differentially expressed transcription factors.

Supplementary Figure 4

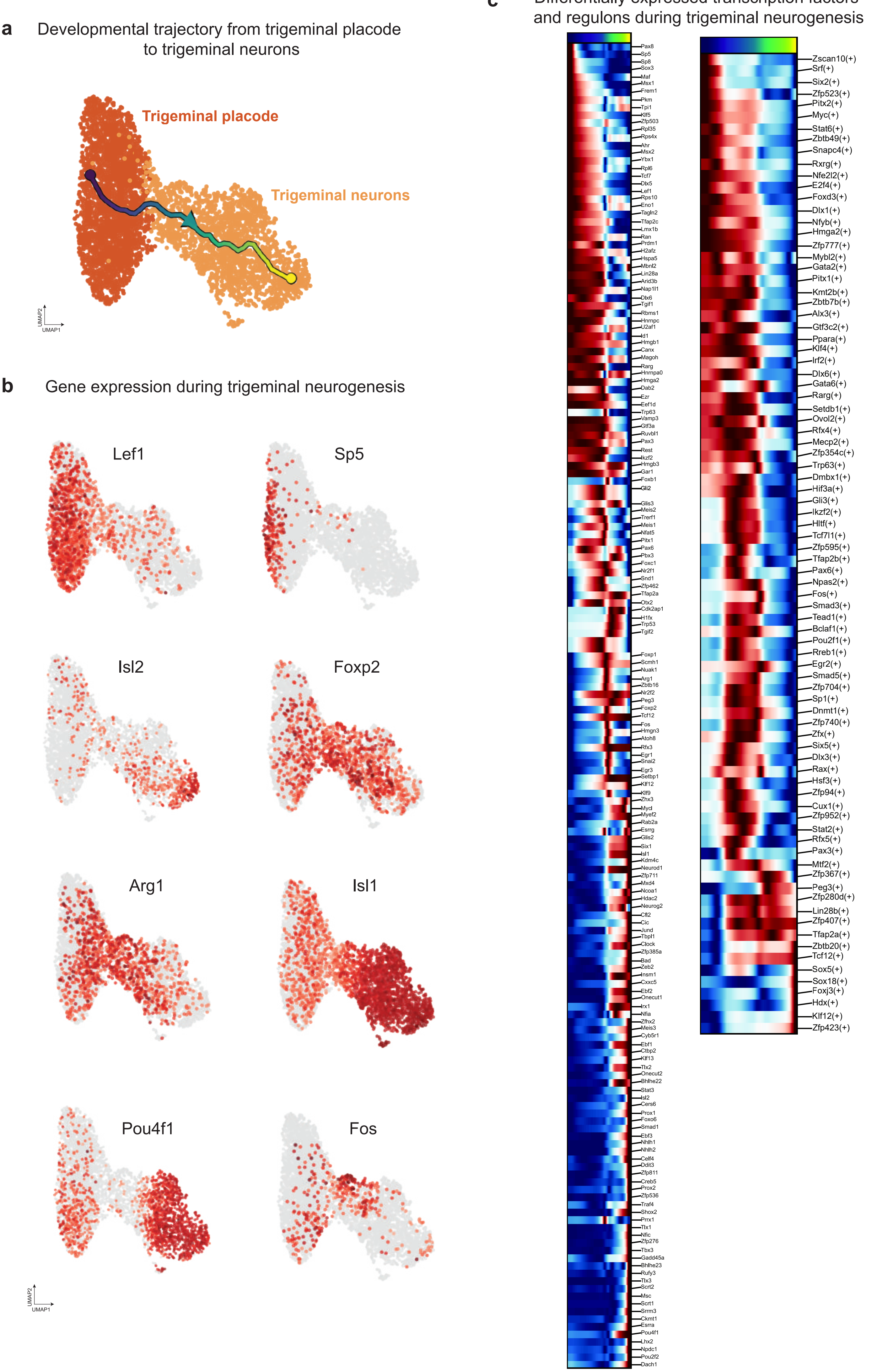

**Supplementary Figure 4. Developmental trajectory from trigeminal placode to trigeminal ganglion.** **(a)** Developmental trajectory from trigeminal placode to trigeminal neurogenesis. **(b)** Visualization of marker gene expression across trigeminal neurogenesis cell states **(c)** Pseudotime-aligned heatmap of differentially expressed transcription factors (left) and active regulons (right) (detected by SCENIC) during trigeminal ganglia development.

#### Supplementary Figure 5

**a** Developmental trajectory from otic placode to otic neurons

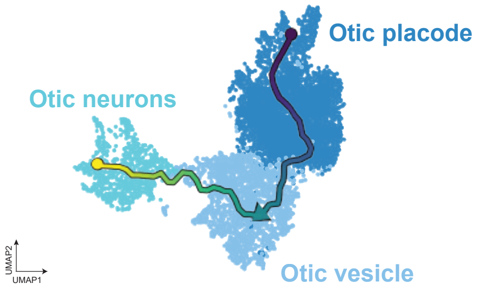


**b** Gene expression during otic neurogenesis

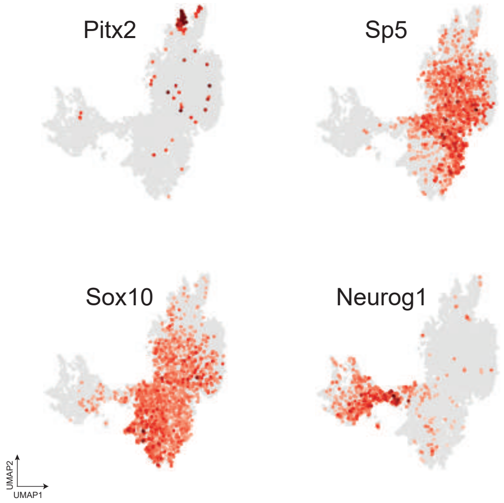


**c** Differentially expressed transcription factors and regulons during otic neurogenesis

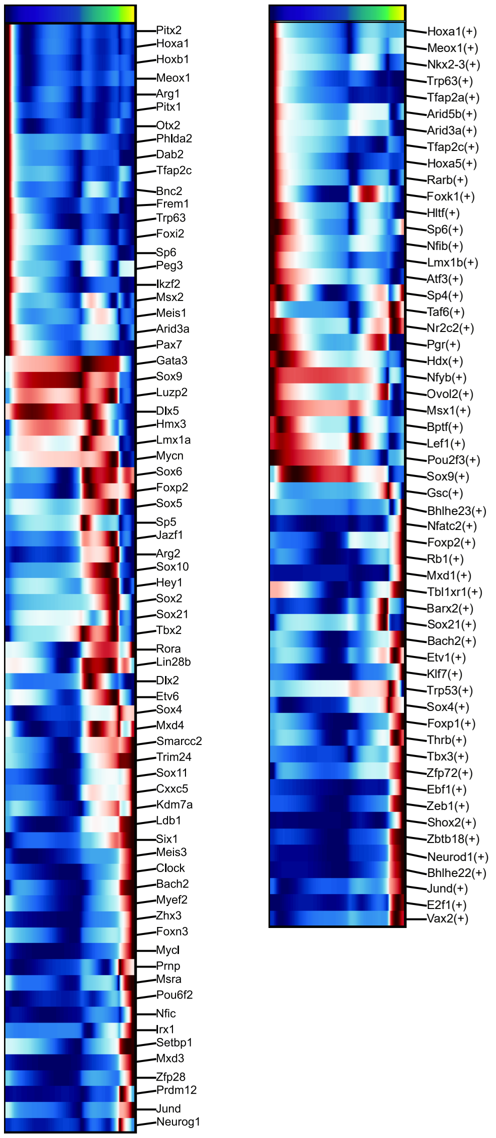

**Supplementary Figure 5. Developmental trajectory from otic placode to cochlear neurogenesis.**
**(a)** Developmental trajectory from otic placode to otic neurogenesis. **(b)** Visualization of marker gene expression across otic neurogenesis cell states **(c)** Pseudotime-aligned heatmap of differentially expressed transcription factors (left) and active regulons (right) (detected by SCENIC) during otic ganglia development.

### Supplementary Figure 6

**a** Developmental trajectory from epibranchial placode to epibranchial neurons

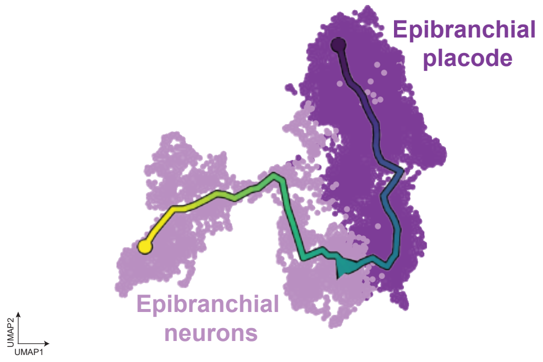


**b** Gene expression during epibranchial neurogenesis

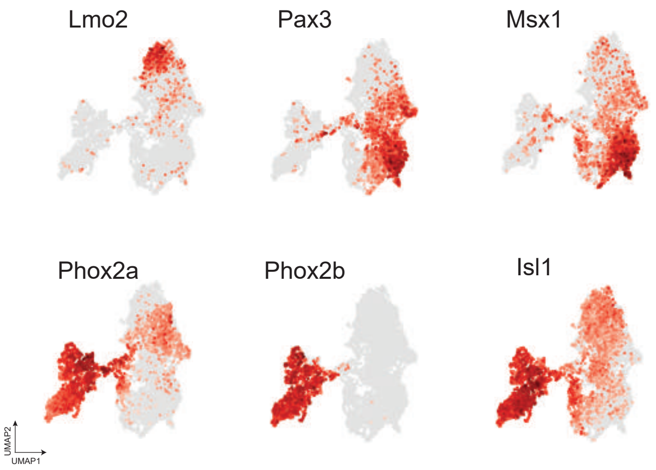


**c** Differentially expressed transcription factors and regulons during epibranchial neurogenesis

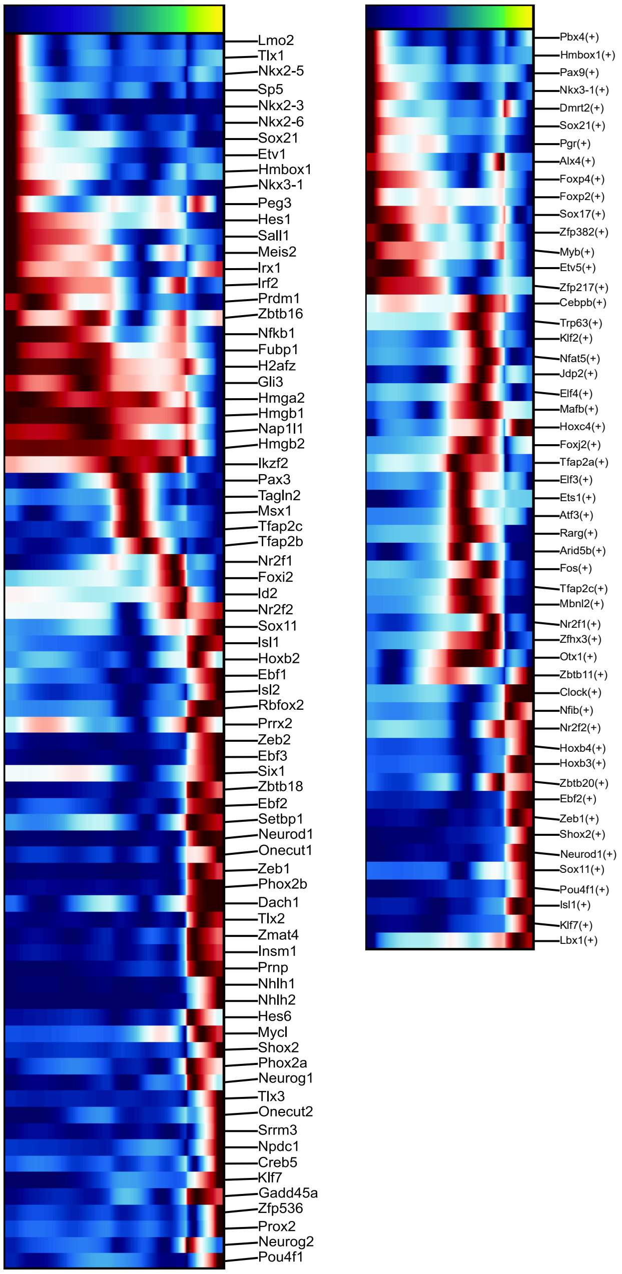

**Supplementary Figure 6. Developmental trajectory from epibranchial placodes to epibranchial ganglia.**

**(a)** Developmental trajectory from epibranchial placodes to neuronal cell populations. **(b)** Visualization of marker gene expression across epibranchial placode and neurogenic cell states **(c)** Pseudotime-aligned heatmap of differentially expressed transcription factors (left)and active regulons (right)(detected by SCENIC) during epibranchial ganglia development.

Supplementary Figure 7

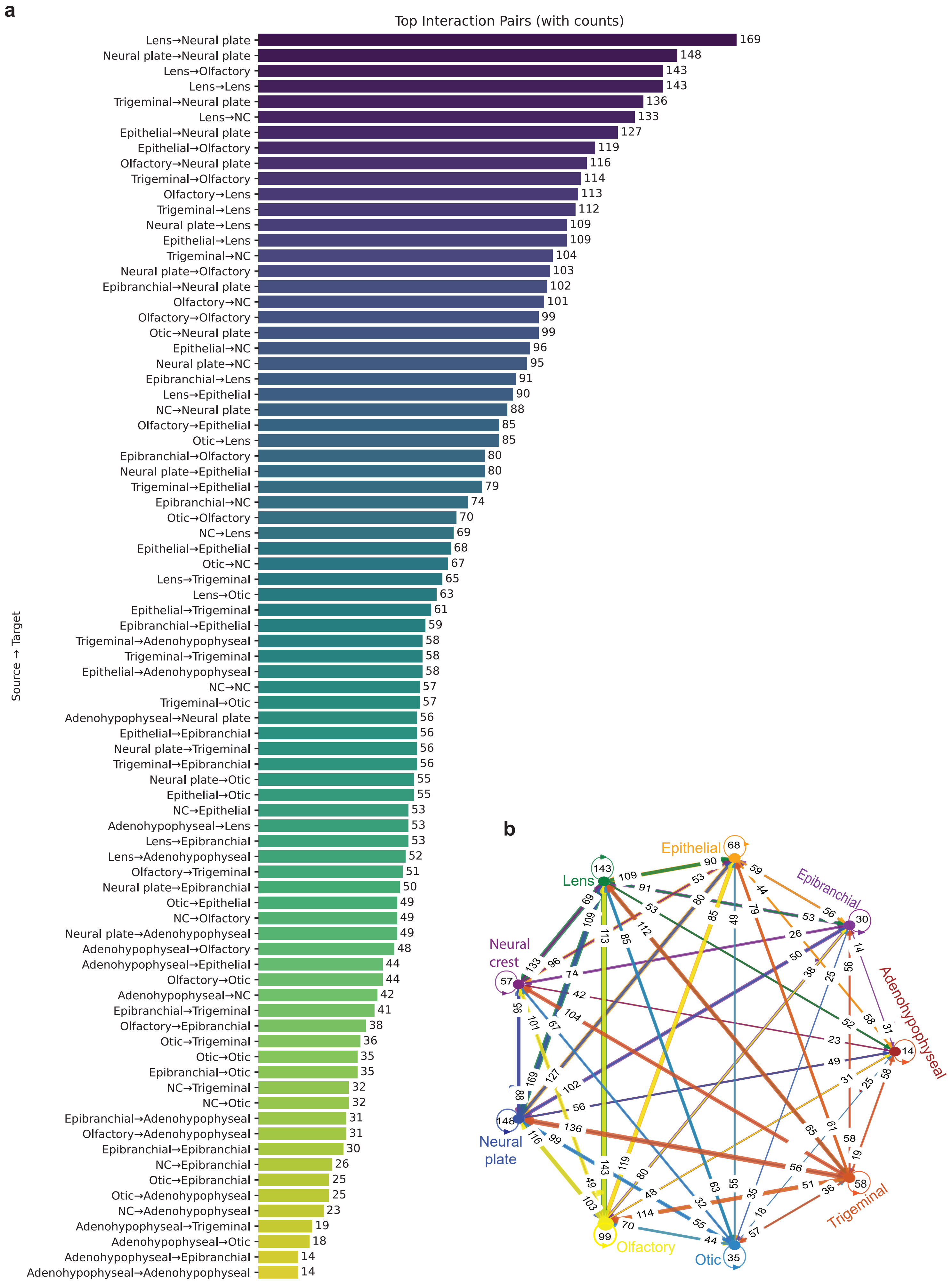

**Supplementary Figure 7. Predicted cell-cell signaling interactions in development of neural plate border derivatives. (a)** Ranking pairwise interactions between different cell subtypes - the higher the bar, the more interactions or stronger the connection. **(b)** A network diagram-visualizing cell types interactions, with color-coded nodes.

Supplementary Figure 8

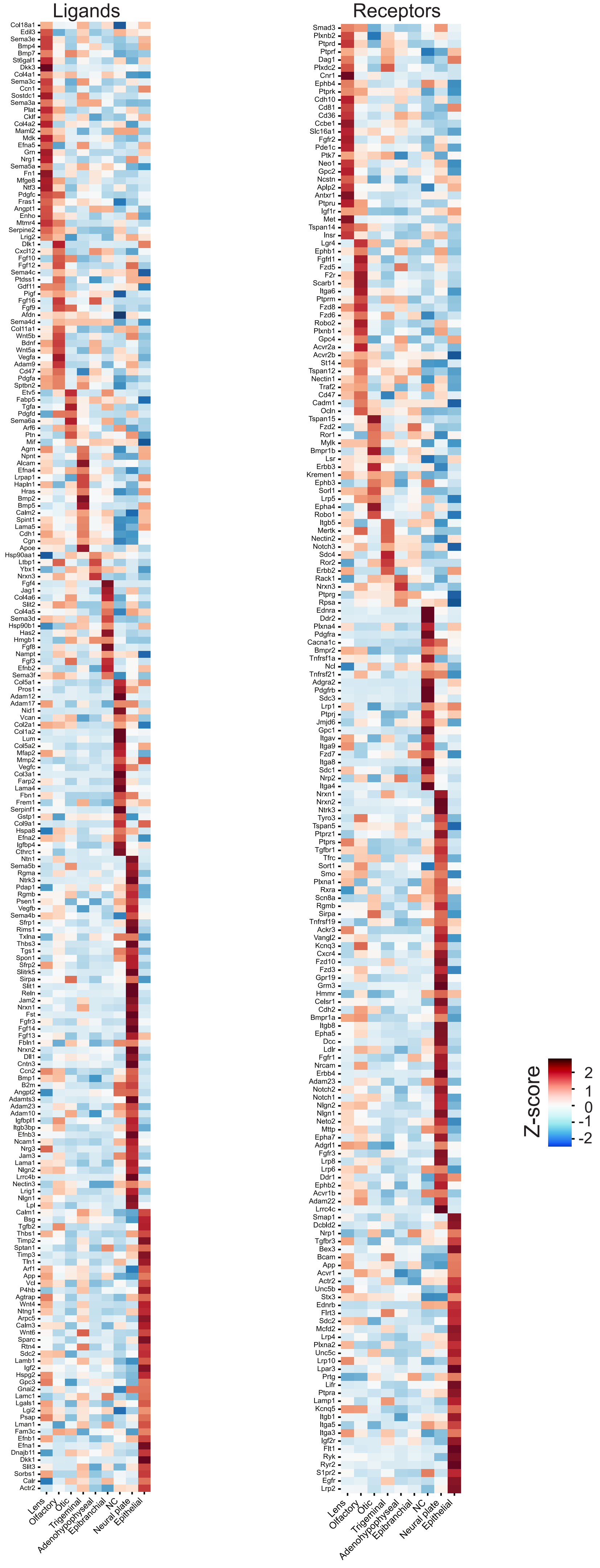

**Supplementary Figure 8. Differentially expressed ligands and receptors in neural plate border derivatives.**

### Supplementary Figure 9

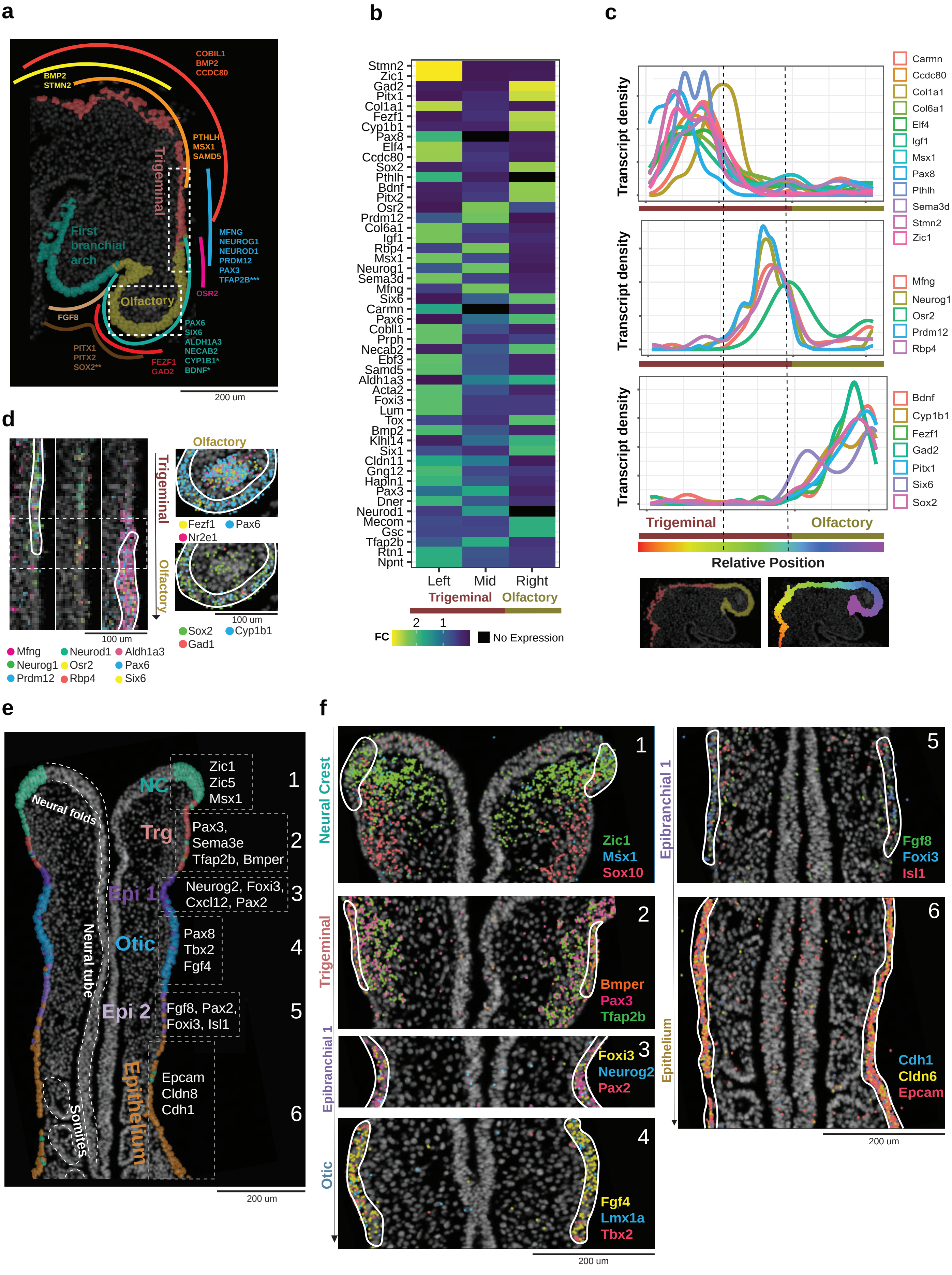

**Supplementary Figure 9. Spatial transcriptomics (Xenium v1) reveals gradual transitions across embryonic ectoderm and neural plate border derivatives, including forming cranial placodes.**
**(a)** Placode-containing and other epithelial clusters together with distribution of highly variable epithelial-specific genes in E8.5 mouse embryo. **(b)** Heatmap showcasing expressed genes over the transition from trigeminal to olfactory placodal clusters within epithelium. **(c)** Line graph showing the expression of genes associated with transition from trigeminal to olfactory clusters over the linearized epithelium. **(d)** Spatial distribution of detected individual transcripts in transitional zone from trigeminal to olfactory regions. **(e)** Spatial distribution of cells assigned to different epithelial clusters including the neural crest, trigeminal, epibranchial, otic and other epithelial domains in an E8.5 mouse embryo. **(f)** Spatial distribution of detected individual transcripts in the neural crest, trigeminal, epibranchial, otic and epithelial regions.

Supplementary Figure 10

**A single paraffin block containing 15 embryos at E8.5, section 1**

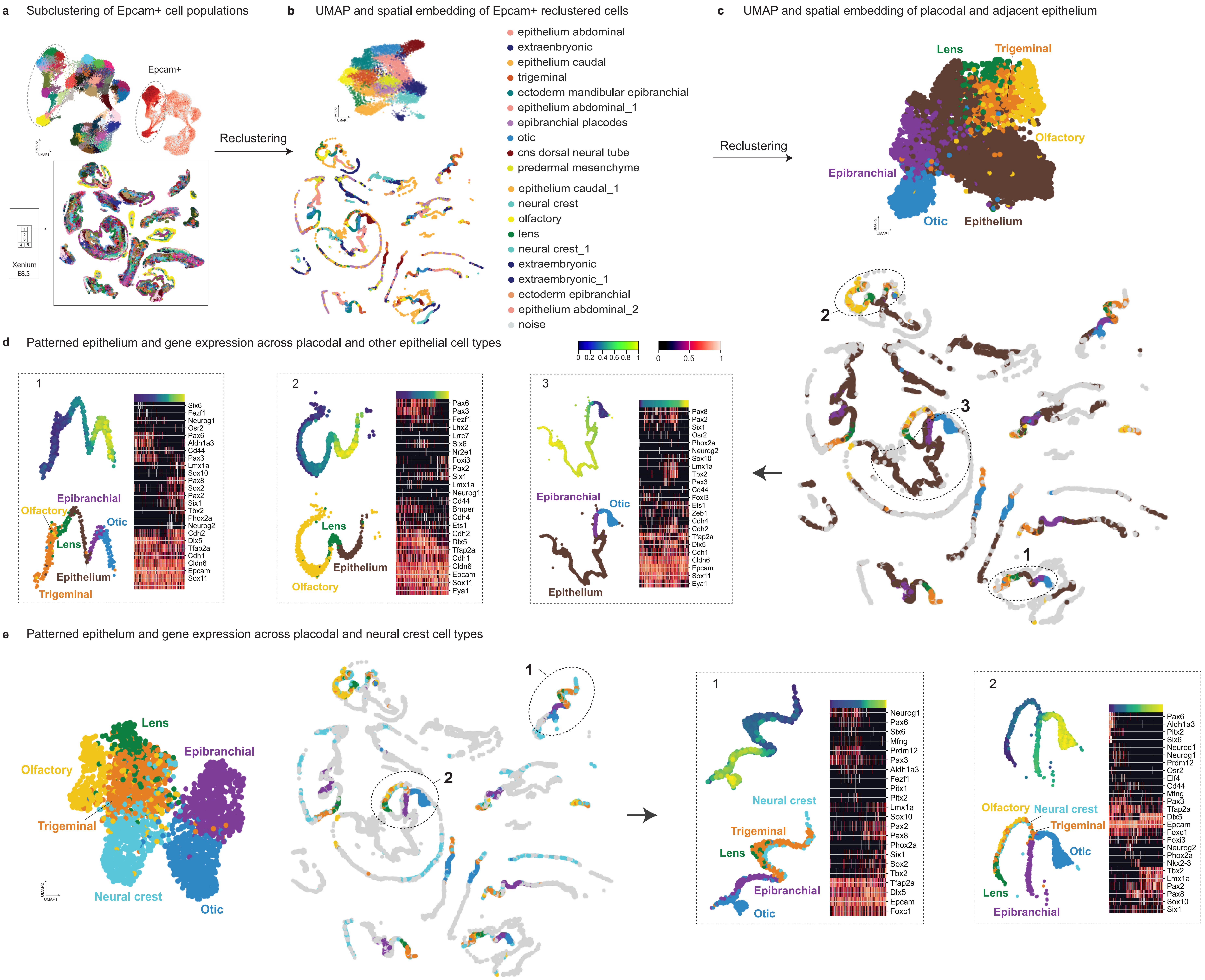

**Supplementary Figure 10. Spatial transcriptomics (Xenium v1) of the neural plate border and other epithelial populations (single paraffin block, E8.5, section 1).**

**(a)** Left, UMAP embedding of the Xenium slide – section 1 at E8.5. Right top, feature plot showing Epcam expression. Right bottom, spatial embedding (segmented epithelial populations). **(b)** UMAP and spatial embedding of *Epcam*+ cells. **(c)** UMAP showing unbiased clusters and the spatial embedding of placode-containing epithelial clusters + non-neural ectoderm. **(d)** Spatial outline of the epithelium, and a gene expression map across placodal and adjacent epithelial regions. Inserts 1, 2, 3, 4, 5 represent heatmaps showing the variable genes over neighboring placodal and other epithelial regions along the spatial coordinates axis. **(e)** UMAP and spatial embedding of placode-containing clusters together with delaminating and migrating cranial neural crest cells. Inserts 1 and 2 contain heatmaps showing the expression of variable genes over the coordinates of exemplified adjacent placodal and neural crest regions.

Supplementary Figure 11

**A single paraffin block containing 15 embryos at E8.5, section 2**

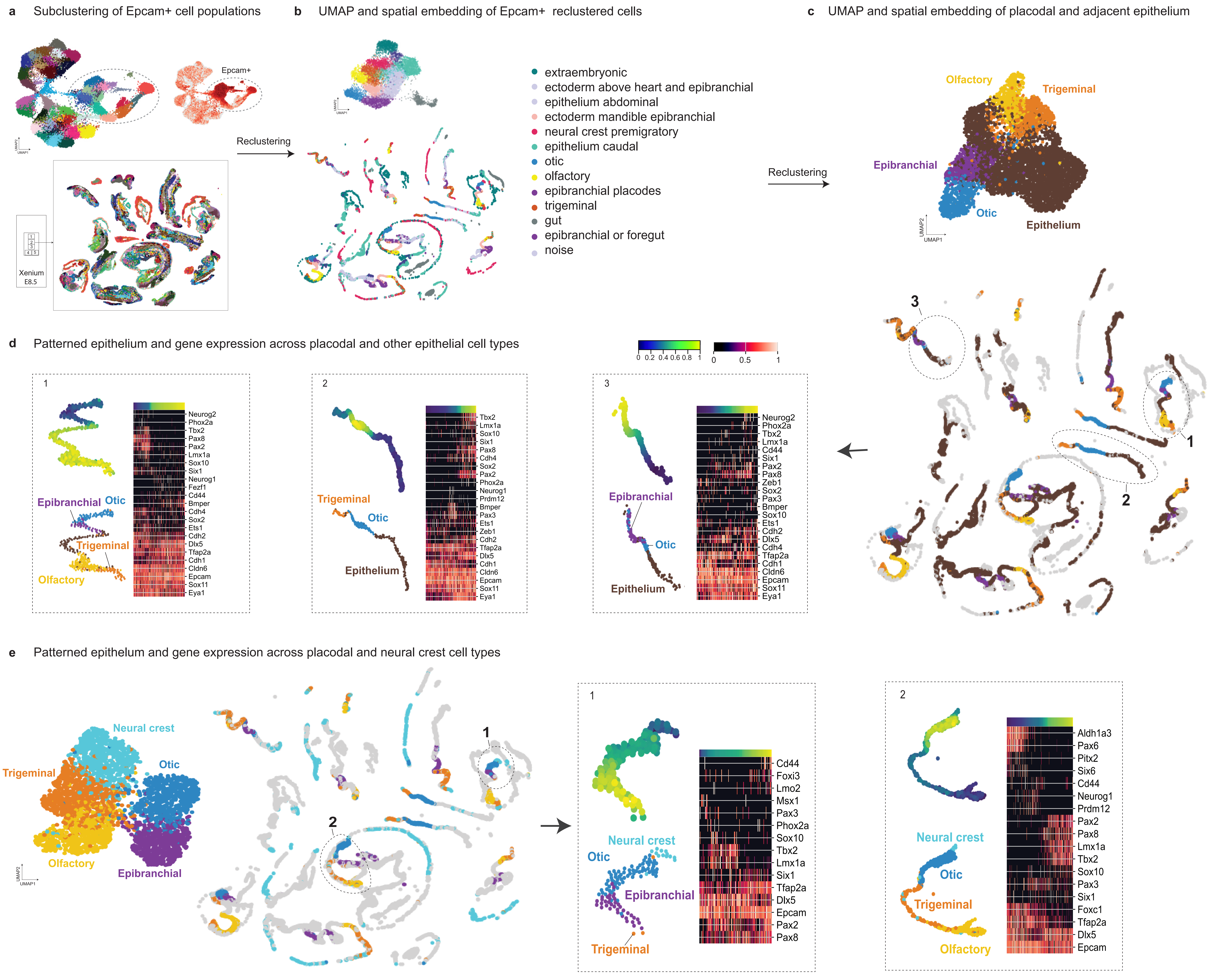

**Supplementary Figure 11. Spatial transcriptomics (Xenium v1) of the neural plate border and other epithelial populations (single paraffin block, E8.5, section 2).**

**(a)** Left, UMAP embedding of the Xenium slide – section 2 at E8.5. Right top, feature plot showing Epcam expression. Right bottom, spatial embedding of identified clusters corresponding to major embryonic cell types. **(b)** Top, UMAP and bottom, spatial embedding of Epcam reclustered cells. **(c)** Left, UMAP and right, spatial embedding of placodal and adjacent epithelial regions. **(d)** Spatial outline of the epithelium, and a gene expression map across placodal and adjacent epithelial regions. Inserts 1, 2, 3, 4 represent heatmaps showing the variable genes over neighboring placodal and other epithelial regions along the spatial coordinates axis. **(e)** UMAP and spatial embedding of placode-containing clusters together with delaminating and migrating cranial neural crest cells. Inserts 1 and 2 contain heatmaps showing the expression of variable genes over the coordinates of exemplified adjacent placodal and neural crest regions.

Supplementary Figure 12

**A single paraffin block containing 13 embryos at E8.5, section 3**

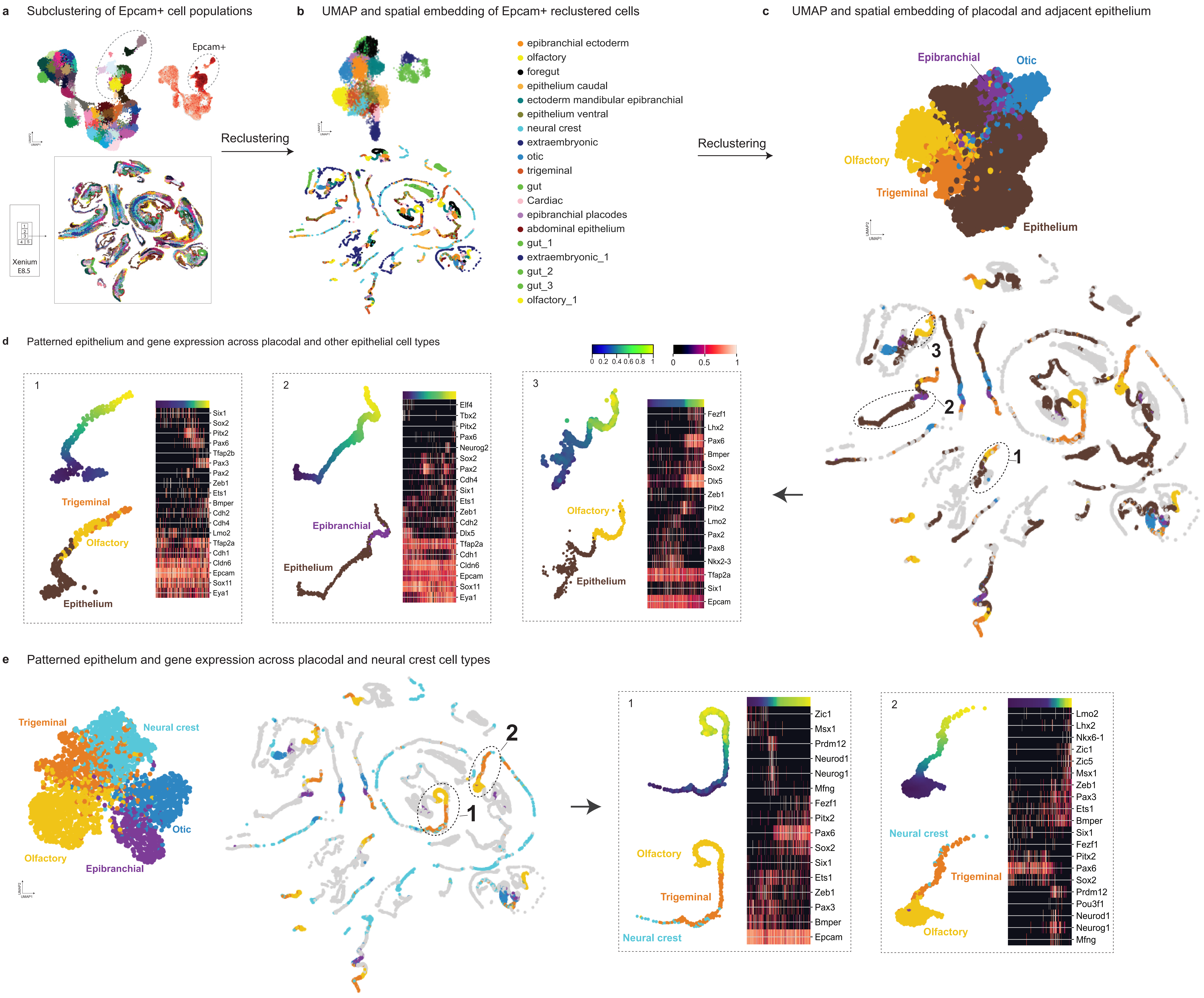

**Supplementary Figure 12. Spatial transcriptomics (Xenium v1) of the neural plate border and other epithelial populations (single paraffin block, E8.5, section 3).**

**(a)** Left, UMAP embedding of the Xenium slide – section 3 at E8.5. Right top, feature plot showing Epcam expression. Right bottom, spatial embedding of identified clusters corresponding to major embryonic cell types. **(b)** Top, UMAP and bottom, spatial embedding of Epcam reclustered cells. **(c)** Left, UMAP and right, spatial embedding of placodal and adjacent epithelial regions. **(d)** Spatial outline of the epithelium, and a gene expression map across placodal and adjacent epithelial regions. Inserts 1, 2, 3, 4 represent heatmaps showing the variable genes over neighboring placodal and other epithelial regions along the spatial coordinates axis. **(e)** UMAP and spatial embedding of placode-containing clusters together with delaminating and migrating cranial neural crest cells. Inserts 1 and 2 contain heatmaps showing the expression of variable genes over the coordinates of exemplified adjacent placodal and neural crest regions.

### Supplementary Figure 13

#### Mapping anterior placode-containing clusters

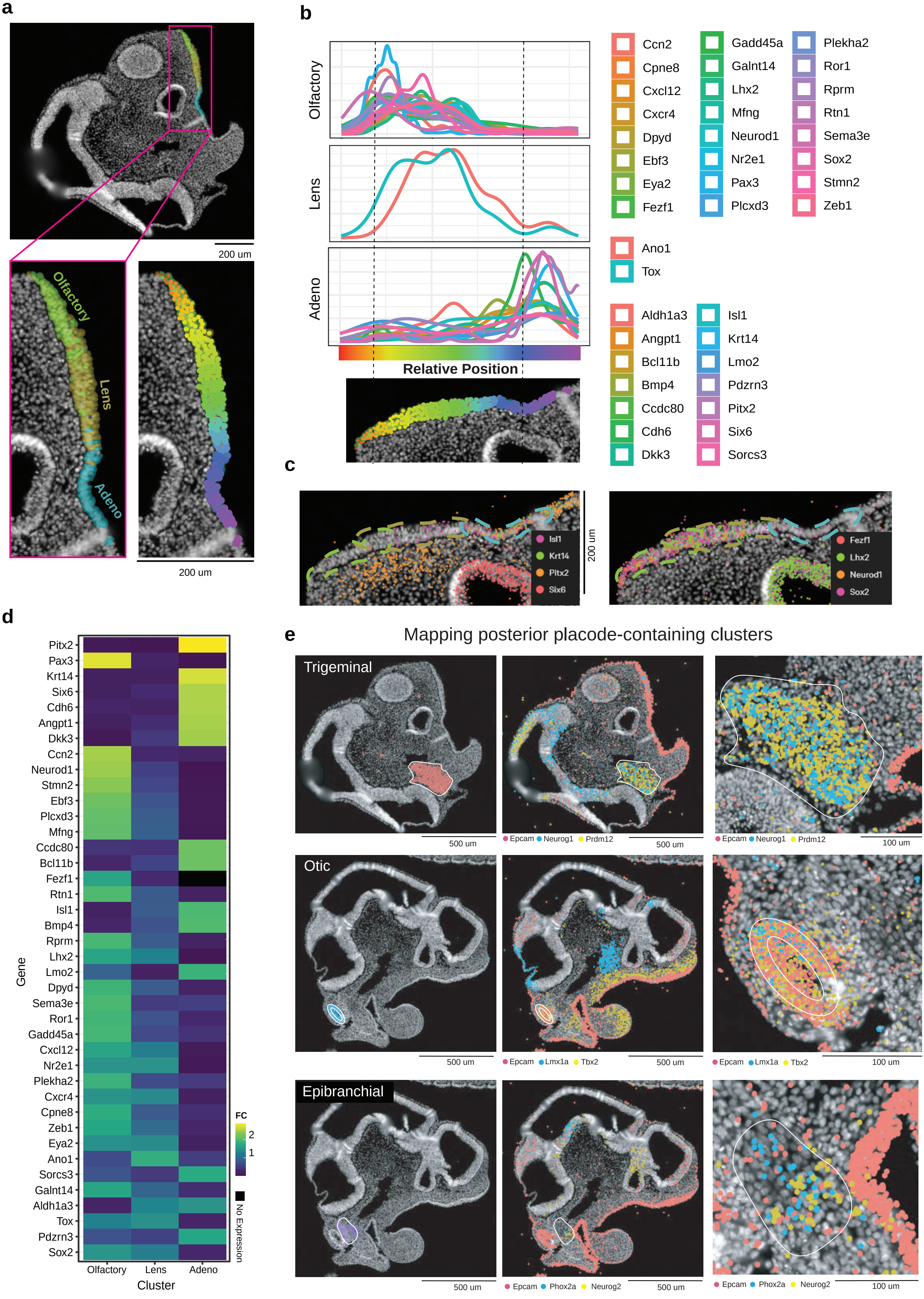

**Supplementary Figure 13. Spatial transcriptomics of the neural plate border derivatives at E9.5.**
**(a)** Spatial transcriptomics representation of annotated cell types in the anterior placodal region (olfactory-, lens-, adenohypophyseal placode-containing clusters) at E9.5. **(b)** Line graphs showing the marker and transitory genes over the linearized epithelium. **(c)** Spatial distributions of actual detected transcripts in the olfactory, adenohypophyseal, and adjacent regions. **(d)** Heatmap showing the characteristic and also transitory expressed genes across the olfactory, lens, and adenohypophyseal placode-containing regions. **(e)** Spatial transcriptomics map of the derivatives of posterior cranial placodes (trigeminal, otic, epibranchial) in E9.5 mouse embryo.

Supplementary Figure 14

a

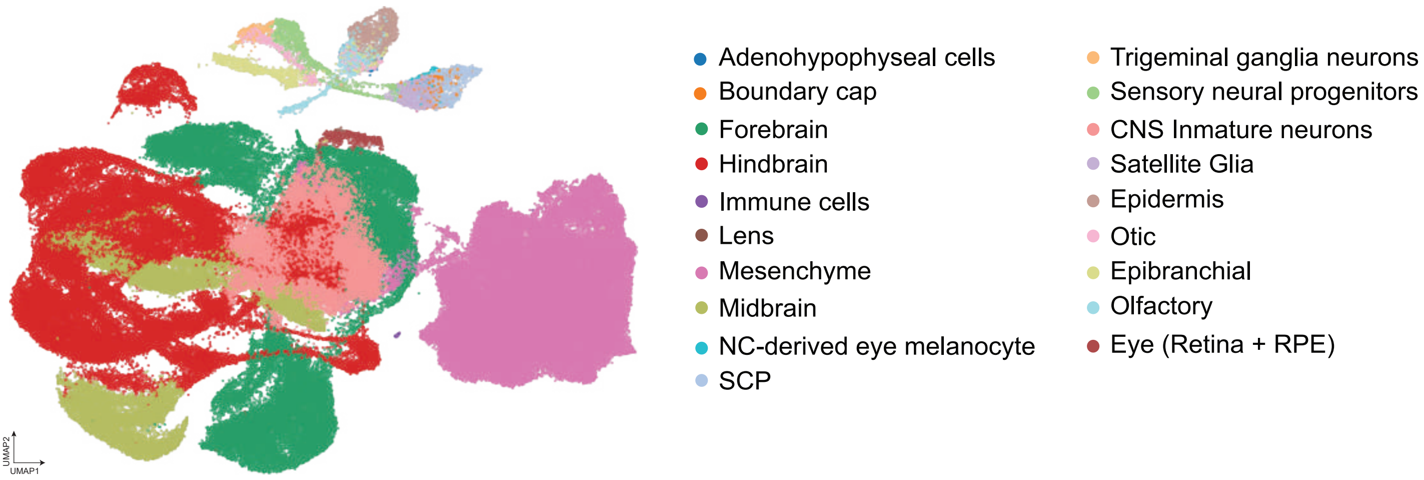


b

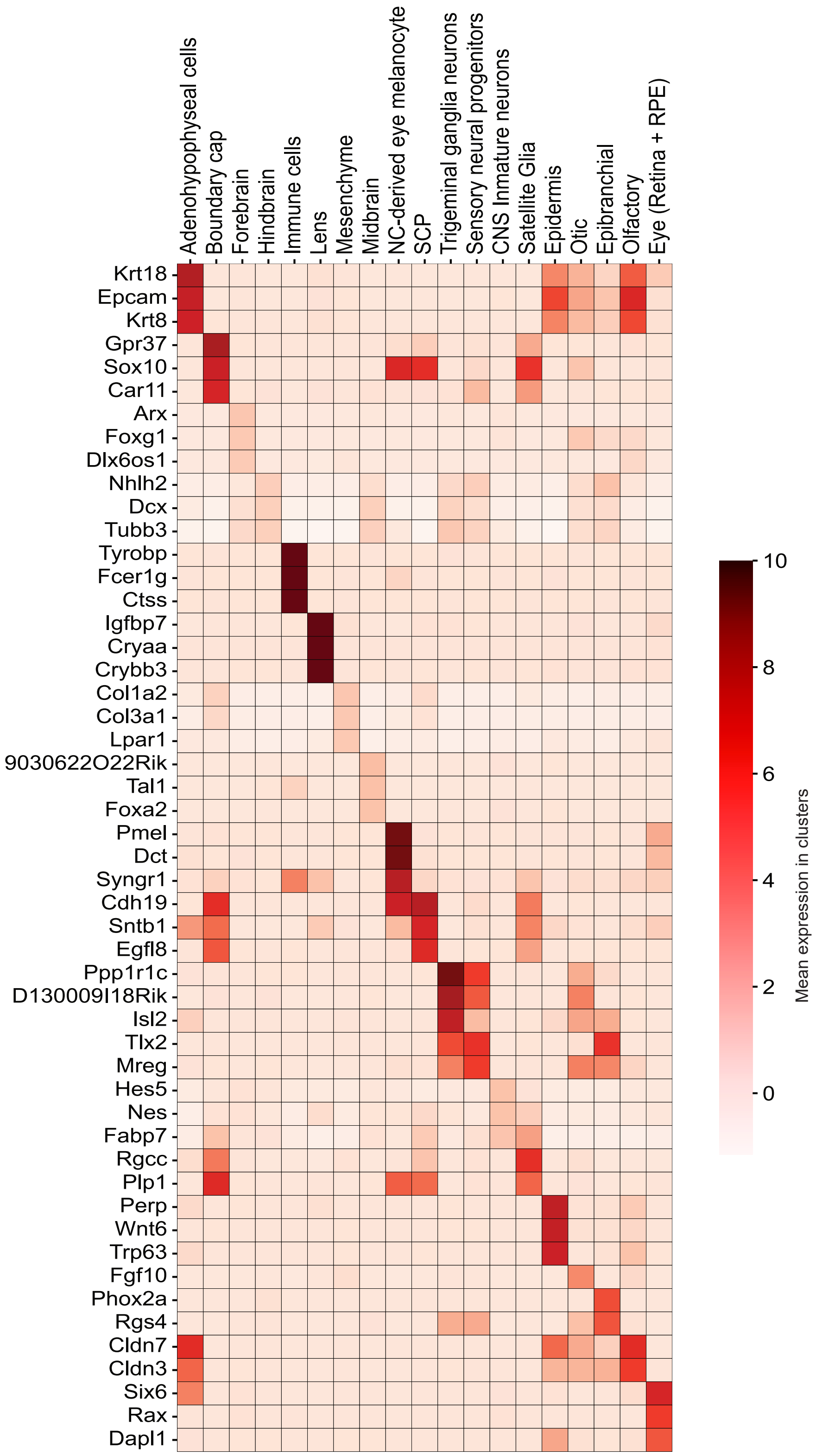

**Supplementary Figure 14. Annotation of clonally barcoded single cell transcriptomics dataset from injections at E7.5-E8).**

**(a)** UMAP embedding showing all major clusters derived from the neural plate border and neural plate. **(b)** Heatmap showing mean expression of selected genes across all annotated clusters.

Supplementary Figure 15

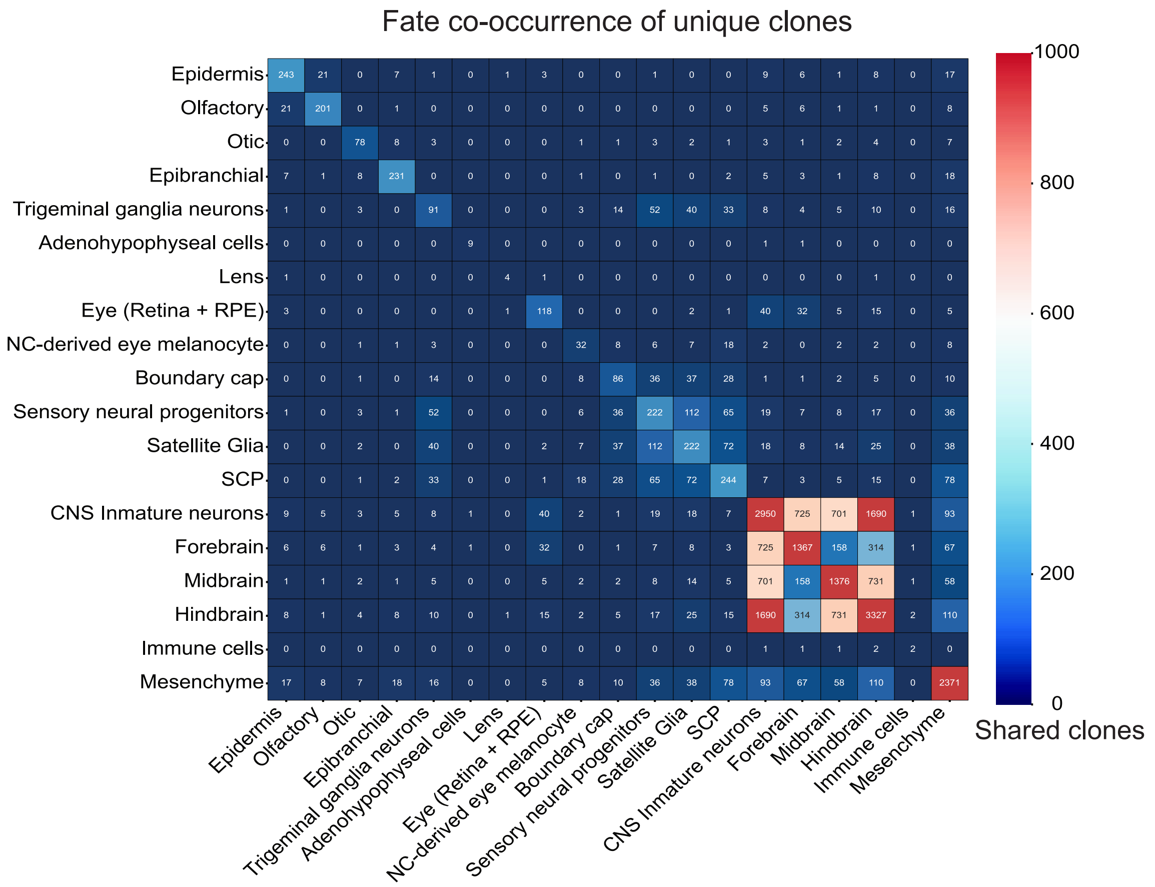

**Supplementary Figure 15. Fate co-occurrence matrix based on clonal relationships.**

Supplementary Figure 16

a

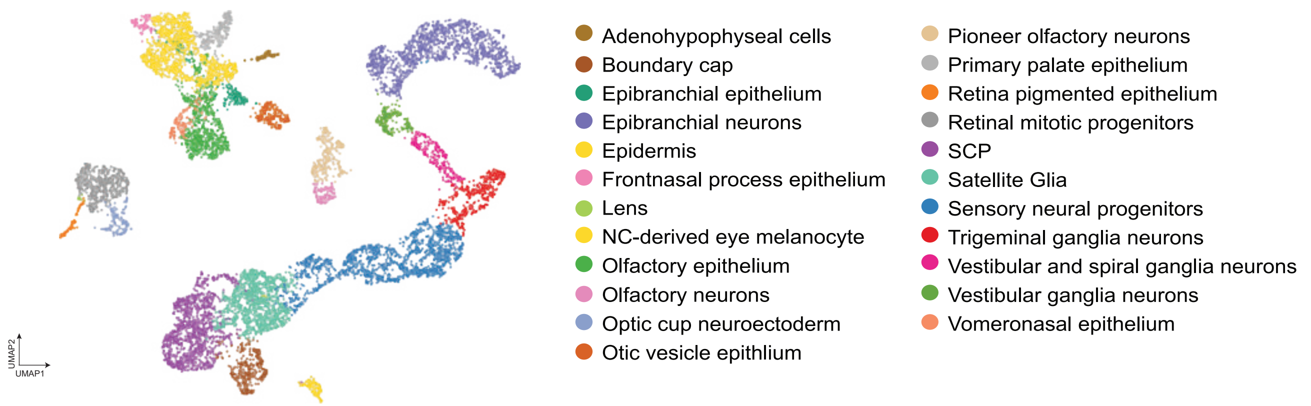


b

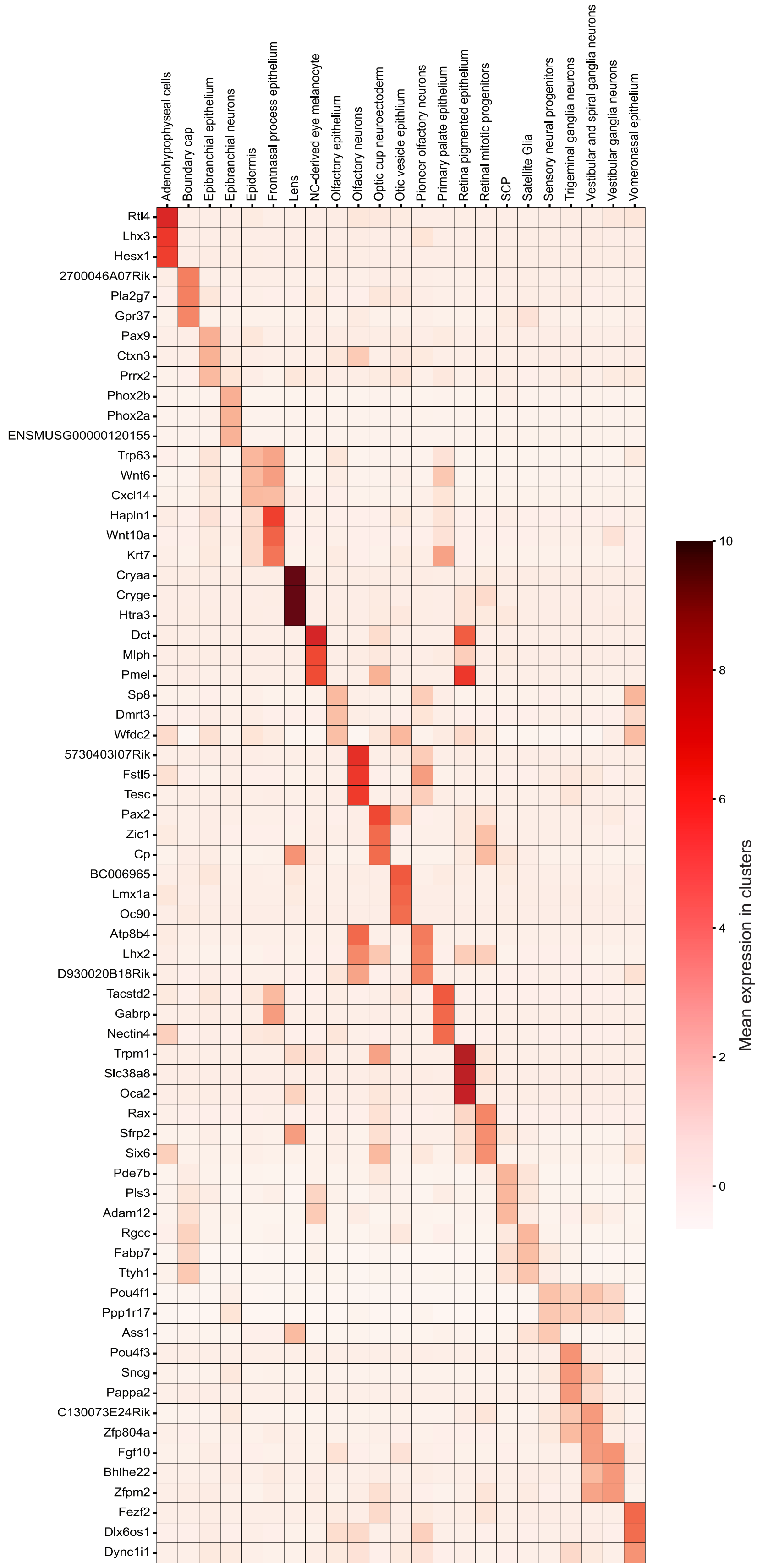

**Supplementary Figure 16. Annotation of clonally barcoded and reclustered placodal and neural crest neuro-glial derivatives.**

**(a)** UMAP of reclustered placodal and neural crest neuro-glial derivatives at E11.5 after injections at E7.5-8. **(b)** Heatmap showing mean expression of selected genes across all annotated clusters.

Supplementary Figure 17

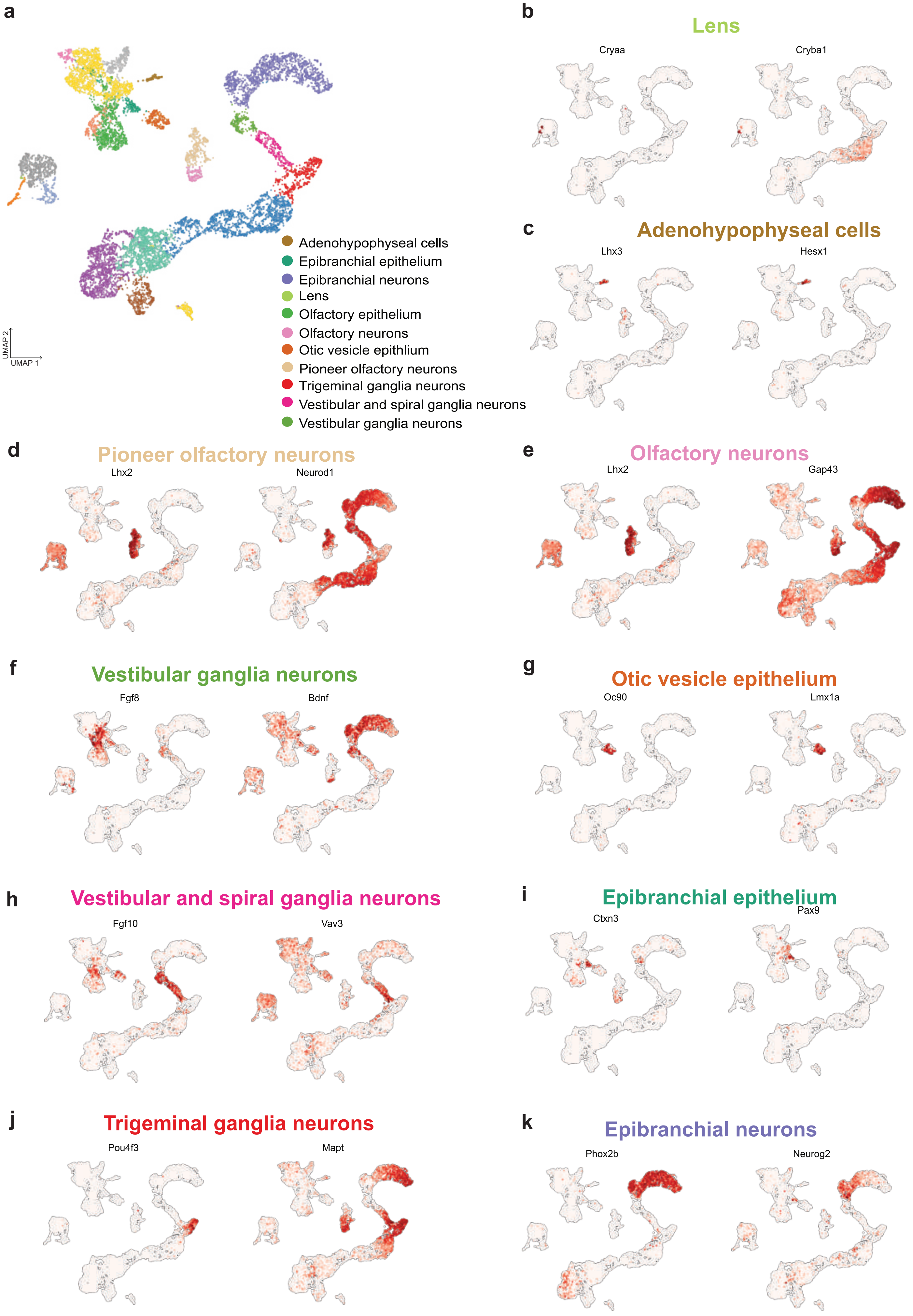

**Supplementary Figure 17. Annotated UMAP and plotted expressed marker genes in clonally barcoded placodal and neural crest neuro-glial derivatives.**
