## Supplementary Table 4 for "Single-cell, clonal and spatial atlases of cranial placodes illuminate their specification and evolution"

**Supplementary Table 4. Quality-control metrics for all single-cell RNA-seq datasets.**

| Sample | Genes by count | Total counts | mt | Doublet rate |
|---|---|---|---|---|
| E8.5 P27217_1001 (wild type) | >2000 | >12000 | < 10 | 0.1% |
| E9.5 P28111 1002 (wild type) | >2000 | >9500 | < 15 | 4.9% |
| E9.5 P28111 1003 (wild type) | >2500 | >8500 | < 14 | 1.3% |
| E9.5 P28111 1004 (wild type) | >1500 | >12000 | < 15 | 0.5% |
| E8.5 P28111 1001 (wild type) | >1000 | >8500 | < 15 | 0.7% |
| E9.5 P29061 1005 (wild type) | >1700 | >3000 | < 15 | 0.2% |
| E8.5 L003 (Foxi3) | >1000 | < 20000 | < 15 | 0.0% |
| E8.5 L004 (Foxi3) | >1000 | | < 15 | 0.1% |
| E8.5 L005 (Foxi3) | < 6000 | < 25000 | < 15 | 0.2% |
| E8.5 L006 (Foxi3) | < 6500 | < 25000 | < 15 | 0.1% |
| E9.5 L003 (Foxi3) | > 1500& < 7000 | > 5000&<140000 | < 15 | 0.3% |
| E9.5 L004 (Foxi3) | > 1500&<6500 | | < 15 | 0.2% |
| E9.5 L005 (Foxi3) | > 1000&<6000 | > 4500 | | 0.2% |
| E9.5 L006 (Foxi3) | > 2000&<6500 | | | 0.3% |
