## Supplementary Table of Tables for "Single-cell, clonal and spatial atlases of cranial placodes illuminate their specification and evolution"

**Supplementary Table 1. Pairwise cell-cell interactions.**

**Supplementary Table 2. Xenium_mBrain_v1.1 metadata and 100 custom genes.**

**Supplementary Table 3. Placode-specific genes and transcription factors.**

**Supplementary Table 4. Quality-control metrics for all single-cell RNA-seq datasets.**
